# Genetic adaptations to SIV across chimpanzee populations

**DOI:** 10.1101/2022.05.24.491624

**Authors:** Harvinder Pawar, Harrison J. Ostridge, Joshua M. Schmidt, Aida M. Andrés

## Abstract

Central and eastern chimpanzees are infected with Simian Immunodeficiency Virus (SIV) in the wild, typically without developing acute immunodeficiency. Yet the recent zoonotic transmission of chimpanzee SIV to humans, which were naïve to the virus, gave rise to the Human Immunodeficiency Virus (HIV), which causes AIDS and is responsible for one of the deadliest pandemics in human history. Chimpanzees have been infected with SIV for tens of thousands of years and have likely evolved to reduce its pathogenicity, becoming semi-natural hosts that largely tolerate the virus. In support of this view, central and eastern chimpanzees show evidence of positive selection in genes involved in SIV/HIV cell entry and immune response to SIV, respectively. We hypothesise that the population first infected by SIV would have experienced the strongest selective pressure to control the lethal potential of zoonotic SIV, and that population genetics will reveal those first critical adaptations. With that aim we used population genomics to investigate signatures of positive selection in the common ancestor of central-eastern chimpanzees. The genes with signatures of positive selection in the ancestral population are significantly enriched in SIV-related genes, especially those involved in the immune response to SIV and those encoding for host genes that physically interact with SIV/HIV (VIPs). This supports a scenario where SIV first infected the central-eastern ancestor and where this population was under strong pressure to adapt to zoonotic SIV. Interestingly, integrating these genes with candidates of positive selection in the two infected subspecies reveals novel patterns of adaptation to SIV. Specifically, we observe evidence of positive selection in numerous steps of the biological pathway responsible for T-helper cell differentiation, including *CD4* and multiple genes that SIV/HIV use to infect and control host cells. This pathway is active only in CD4+ cells which SIV/HIV infects, and it plays a crucial role in shaping the immune response so it can efficiently control the virus. Our results confirm the importance of SIV as a selective factor, identify specific genetic changes that may have allowed our closest living relatives to reduce SIV’s pathogenicity, and demonstrate the potential of population genomics to reveal the evolutionary mechanisms used by naïve hosts to reduce the pathogenicity of zoonotic pathogens.

**Author Summary:** Chimpanzees are at the origin of HIV-1, a virus that generates an incurable disease and that generated a pandemic that has claimed 35 million lives. Chimpanzees have evolved to control the pathogenicity of the virus, which does not typically develop into AIDS in the same way as in humans. Identifying the genetic adaptations responsible for this process provides critical knowledge about SIV and HIV. Our analysis of chimpanzee genetic adaptations identified specific genes and molecular pathways involved in adaptation to SIV, providing important insights into the mechanisms that likely allowed our closest living relatives to control SIV/HIV. Further, we establish SIV as a strong and recurrent selective pressure in central and eastern chimpanzees, two important subspecies of large mammals that are currently endangered.

## Introduction

A typical evolutionary outcome of pathogen infections is the development of host immunity or tolerance to an infectious agent that was initially highly pathogenic, and that remains so in naïve species. Host genetic adaptations underlie the sometimes striking differences in infection outcome in different species infected by the same virus. The lentivirus Simian Immunodeficiency Virus (SIV) provides a fascinating example of this process. Most African primates, the great apes being notable exceptions, have their own endemic SIV strain for which they are considered natural hosts (Chahroudi et al. 2012). When natural hosts (e.g. vervet monkeys) are exposed to their endemic SIV, they have a proportionate immune activation that preserves CD4+ T cell counts and wider immune function, resulting in asymptomatic outcomes and no significant reduction in lifespan despite viral replication (Ma et al. 2013; Ma et al. 2014; Silvestri et al. 2007). In contrast, species that are naïve to SIV, such as non-African primates, are unable to resolve the acute phase of infection and instead generate chronic immune activation, CD4+ T cell depletion and eventual immunodeficiency (Goldstein et al. 2005; Mandell et al. 2014; Barouch et al. 2016; Chahroudi et al. 2012). The course of these infections and the development of clinical disease is almost identical to that of humans infected with HIV, which are also a naïve species and which, without treatment, progress to acquired immunodeficiency syndrome (AIDS).

Humans have received multiple zoonotic transmissions of SIV. The most deadly, which introduced HIV-1 group M responsible for the AIDS pandemic, originated when the SIV of a central chimpanzee (*Pan troglodytes troglodytes*) jumped into humans in the early twentieth century (Keele et al. 2006; Worobey et al. 2008). HIV-1 is thus genetically very similar to chimpanzee SIV (SIVcpz) (Keele et al. 2006). Also, the chimpanzee genome has a high sequence identity to the human genome (Chimpanzee Sequencing and Analysis Consortium 2005) and the two species share most aspects of their physiology. Therefore, the mechanisms used by chimpanzees to limit the pathogenicity of SIV may be informative about potential mechanisms to control HIV.

Chimpanzees have traditionally been considered natural SIV hosts, because despite infections in zoos and laboratories, infected chimpanzees rarely progress to AIDS-like symptoms (Sharp et al. 2005; de Groot et al. 2017; Gilden et al. 1986). Further, SIV-infected chimpanzees show proportionate patterns of immune activation, similar to those observed in natural hosts, such as vervet monkeys (Greenwood et al. 2015). Yet, recent observations have suggested that chimpanzees are not true natural hosts, instead lying between natural and symptomatic SIV hosts. Disease symptoms such as CD4+ T-cell depletion and thrombocytopenia have been observed in infected chimpanzees and associated with reduced fitness, particularly in the wild (Keele et al. 2009; Greenwood et al. 2015; Etienne et al. 2011). Specifically, Keele *et al*., observed in habituated wild eastern chimpanzees that SIV infection increased mortality risk and infected females had fewer offspring and higher infant mortality rates than uninfected females (Keele et al. 2009). Similarly, SIV infection has been associated with population decline in the Kalande community of eastern chimpanzees (Rudicell et al. 2010). It should be noted that studies investigating the fitness effects of SIVcpz in the wild have been restricted to small sample sizes from few habituated populations due to the inherent challenges associated with studying wild populations of endangered large mammals. Nevertheless, evidence suggests that chimpanzees can be considered ‘semi-natural’ hosts, as the virus lacks the dramatic pathogenicity seen in naïve species like humans yet likely has non-negligible health consequences and fitness effects. This is perhaps expected if chimpanzees have already acquired adaptations that control the high initial pathogenicity of SIV but have not yet acquired adaptations that fully control the effects of the infection.

Interestingly, of the four genetically and geographically distinct subspecies of chimpanzee, natural SIV infection is found only in central (*P. t. troglodytes*) and eastern (*P. t. schweinfurthii*) chimpanzees, who are most closely related to each other (Prado-Martinez et al. 2013) (Figure 1). SIV infection has not been detected in the wild in the second clade of chimpanzees, which includes the western (*P. t. verus*) and Nigeria-Cameroon (*P. t. ellioti*) subspecies (Locatelli et al. 2016; Keele et al. 2006; Prado-Martinez et al. 2013). The uneven distribution of SIV among chimpanzee subspecies has sparked much interest in the origin of the virus in the *Pan* lineage.

**Figure 1.**
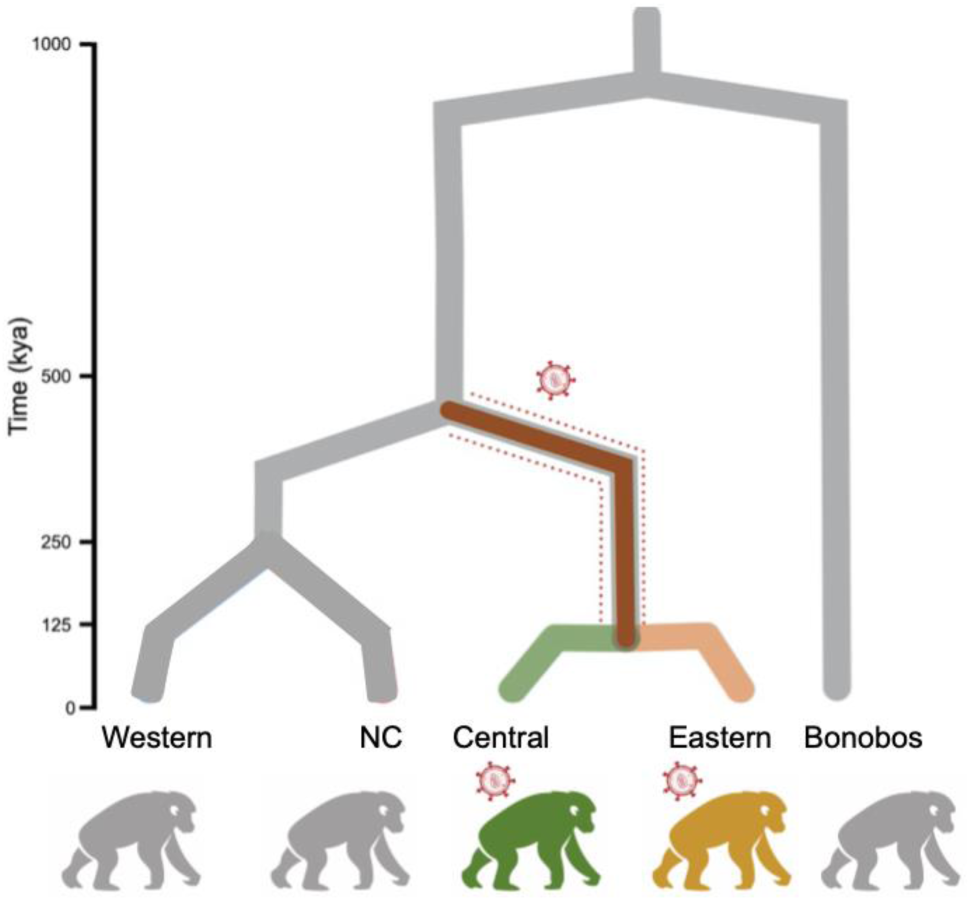
Phylogeny of chimpanzee subspecies and bonobos with the distribution of natural SIV infection. The dotted red line indicates the central-eastern ancestor where the first SIV infection most likely occurred, for which we investigate positive selection.

We know that SIVcpz is the result of zoonotic transmission to chimpanzees of at least two African monkey SIV strains (SIV from the red-capped mangabey and the ancestor of the SIVs currently infecting mona, moustached and greater spot-nosed monkeys), which then recombined and generated SIVcpz, a mosaic lineage able to infect and transmit in chimpanzees (Bell and Bedford 2017; Bailes et al. 2003). Given this complex origin, it is most likely that SIVcpz arose only once. More than one scenario is theoretically compatible with the distribution of SIVcpz in nature. Zoonotic transmission to the common ancestor of all chimpanzees and subsequent clearance in western and Nigeria-Cameroon is highly unlikely given the current SIVcpz distribution (Sharp, Shaw, and Hahn 2005; Santiago et al. 2002), as it would require the virus to be lost in two subspecies. Zoonotic transmission into the central-eastern clade is more likely, either through transmission to the common ancestor of central and eastern chimpanzees or through transmission to one subspecies (e.g. central) with subsequent transfer to the other subspecies (e.g. eastern). A reliable estimate of the time of the most recent common ancestor of central and eastern SIVcpz could be informative about the likelihood of these two scenarios, but the imprecision of phylogenetic dating of SIVcpz, which has been highly variable (Wertheim and Worobey 2009; Worobey et al. 2010) due to the difficulties associated with dating deep times scales for SIV (Worobey et al. 2010), complicate such inferences. The phylogeny of SIVcpz separates two monophyletic sister clades, one containing all SIVcpzPtt sequences (the virus infecting central chimpanzees) and one containing all SIVcpzPts sequences (the virus infecting eastern chimpanzees) (Leitner et al. 2007; Sharp, Shaw, and Hahn 2005), which could be compatible with both scenarios. SIVcpzPts lineages do not fall within the diversity of SIVcpzPtt, or vice versa (Leitner et al. 2007), as we would expect if one subspecies (e.g. centrals) was originally infected and recently infected the other subspecies (e.g. easterns). However, if SIVcpz was laterally transmitted sufficiently long ago, lineage extinction could generate reciprocal monophylogeny in this scenario too. So while the two scenarios are not equally likely, distinguishing among them is difficult with current data.

SIV is a dangerous pathogen, and as such a strong selective force. Once SIV infects a species, SIV-related adaptation is likely to be pervasive and continuous. In fact even natural hosts such as vervet monkeys, which have been infected for 0.5-3 million years, show genome-wide signatures of positive selection in SIV-related genes (Svardal et al. 2017; Ma et al. 2014). It has long been thought that SIV may be a strong selective force in chimpanzees, posited to drive allele frequency change and adaptation in a few immune and SIV-related genes (de Groot et al. 2002; de Groot et al. 2000; de Groot et al. 2008; Wooding et al. 2005; MacFie et al. 2009). Analysing dozens of genomes, we confirmed that SIV has driven chimpanzee evolution by uncovering the presence of recent adaptations to SIV in the central and eastern subspecies (Schmidt et al. 2019). Using the PBSnj statistic, we identified the SNPs with the greatest allele frequency change in each chimpanzee subspecies, which are the strongest candidate targets of subspecies-specific positive selection. Strikingly, both in central and eastern chimpanzees, these SNPs are enriched in genes related to SIV. As expected under a model of zoonotic transmission into the central-eastern clade, neither western nor Nigeria-Cameroon showed evidence of recent positive selection in SIV-related gene categories (Schmidt et al. 2019). Although earlier work had reported evidence of low diversity at *CCR5, CXCR4* and *CX3CR1* in Nigeria-Cameroon and western, and suggested that this may be due to positive selection (Wooding et al. 2005; MacFie et al. 2009), it is unclear if SIV was the selective force.

The evidence of genetic adaptation in sets of SIV-related genes is clearer and easy to interpret. Identified candidate targets of recent selection in central chimpanzees are enriched in cytokine coreceptors due to signatures in *CCR3*, *CCR9* and *CXCR6* (Schmidt et al. 2019), which mediate HIV cell entry together with the primary receptor *CD4* (Elliott et al. 2015; Wetzel et al. 2017; Nedellec et al. 2009) and are paralogs of the HIV coreceptors *CCR5* and *CXCR4* (Berger 1997; Moore et al. 2004; Blumenthal et al. 2012). In contrast, selection targets in eastern chimpanzees are enriched in “SIV-response genes”, genes that upon SIV infection show different expression profiles in CD4+ T lymphocytes in natural host vs. naïve host species (in this case, vervet monkeys vs macaques, (Svardal et al. 2017; Jacquelin et al. 2014; Jacquelin et al. 2009)). These SIV-response genes likely contribute to the finely tuned natural host response to SIV, which is able to control the infection and results in non-pathogenic outcomes (Schmidt et al. 2019). Thus, interestingly, the two subspecies seem to have evolved differential adaptation mechanisms to control SIV.

Under a scenario of zoonotic transmission into the central-eastern ancestor, this naïve population would have been under strong pressure to adapt –actually, it would be the chimpanzee population with the strongest pressure to adapt. We thus hypothesise that under this scenario positive selection in the central-eastern ancestor would have been critical to control the lethal potential of zoonotic SIV upon a naïve chimpanzee population. Under that hypothesis, we could detect signatures of positive selection in SIV-related genes in the central-eastern ancestor. We test this hypothesis using 47 chimpanzee whole genomes. We find evidence of adaptation in genes involved in SIV biology, providing further support for SIV infection in the central-eastern ancestor. Further, candidate targets of positive selection point to diverse adaptive mechanisms, including host response to infection, viral interactions and cell entry. Combining information across populations we discover previously known mechanisms of adaptation to SIV. Specifically, we identify the T helper cell type-1/type-2 (Th1/Th2) differentiation pathway as a critical player that has been repeatedly targeted by positive selection at different time points during chimpanzee evolution. This reveals the potential of this functional pathway, and specific genes within the pathway, to control the pathogenicity of SIV/HIV infection.

Excitingly, population genetics tools allow us to identify the genetic adaptations that happened at this critical time during which chimpanzees evolved to reduce the pathogenicity of a potentially deadly virus.

## Results

### Signatures of positive selection in the central-eastern ancestor

We identified genomic regions showing evidence of positive selection in the central-eastern ancestor using 3P-CLR (Racimo 2016). 3P-CLR tests distortions of the site frequency spectrum (SFS) due to selective sweeps, by modelling the evolutionary trajectory of alleles in a 3-population tree and comparing the likelihood of the observed SFS under contrasting hypotheses of neutrality and positive selection (Racimo 2016). 3P-CLR is ideal for our purpose. First, it has high power to detect the events we target, as it was designed to identify hard sweeps in ancestral modern humans using Neanderthals as an outgroup (Racimo 2016), and the modern human-Neanderthal divergence (Prüfer et al. 2017) is on the order of the inferred split time between chimpanzee clades (de Manuel et al. 2016). We established with simulations that 3P-CLR also has very high power to detect selective events in the central-eastern ancestor when Nigeria-Cameroon is used as outgroup (Figure S1), with power equal or higher than that reported in humans (Racimo 2016) under the same scenario: strong selection and fixation of the advantageous allele (Supplementary Material). Second, 3P-CLR has high power to detect selection in the ancestral population while being largely unaffected by convergent evolution (independent selection in both central and eastern chimpanzees) that does not generate the SFS distortions that translate into high 3P-CLR likelihood ratio scores (Racimo 2016).

Informed by the power analysis, we applied 3P-CLR to the central–eastern–Nigeria-Cameroon 3-population tree, using the genotypes from high-coverage autosomal genomes of 47 chimpanzees (18 central, 19 eastern and 10 Nigeria-Cameroon) (de Manuel et al. 2016), sliding windows of size 0.25 centiMorgans (cM) and the recombination map of Auton *et al*., (Auton et al. 2012). The windows with the highest 3P-CLR scores in the genome-wide empirical distribution have the strongest evidence of positive selection at this time depth. We thus consider candidate targets of positive selection the windows with the highest 99.5%, 99.9% and 99.95% 3P-CLR likelihood ratio scores in the genome, which correspond to the 0.5% (n=4090), 0.1% (n=818) and 0.05% (n=409) tails of the empirical distribution. The candidate windows have additional signatures that, while not fully independent from the 3P-CLR signatures, are expected under positive selection in the central-eastern ancestor: they contain a marked excess of sites with high derived allele frequencies (DAF) in central and eastern (Figure 2A), and an excess of highly differentiated SNPs between the central-eastern clade and Nigeria-Cameroon (Figure 2B). While we cannot discard the presence of some false positives, these genomic regions are prime candidates to have mediated genetic adaptations in the central-eastern ancestral population.

**Figure 2.**
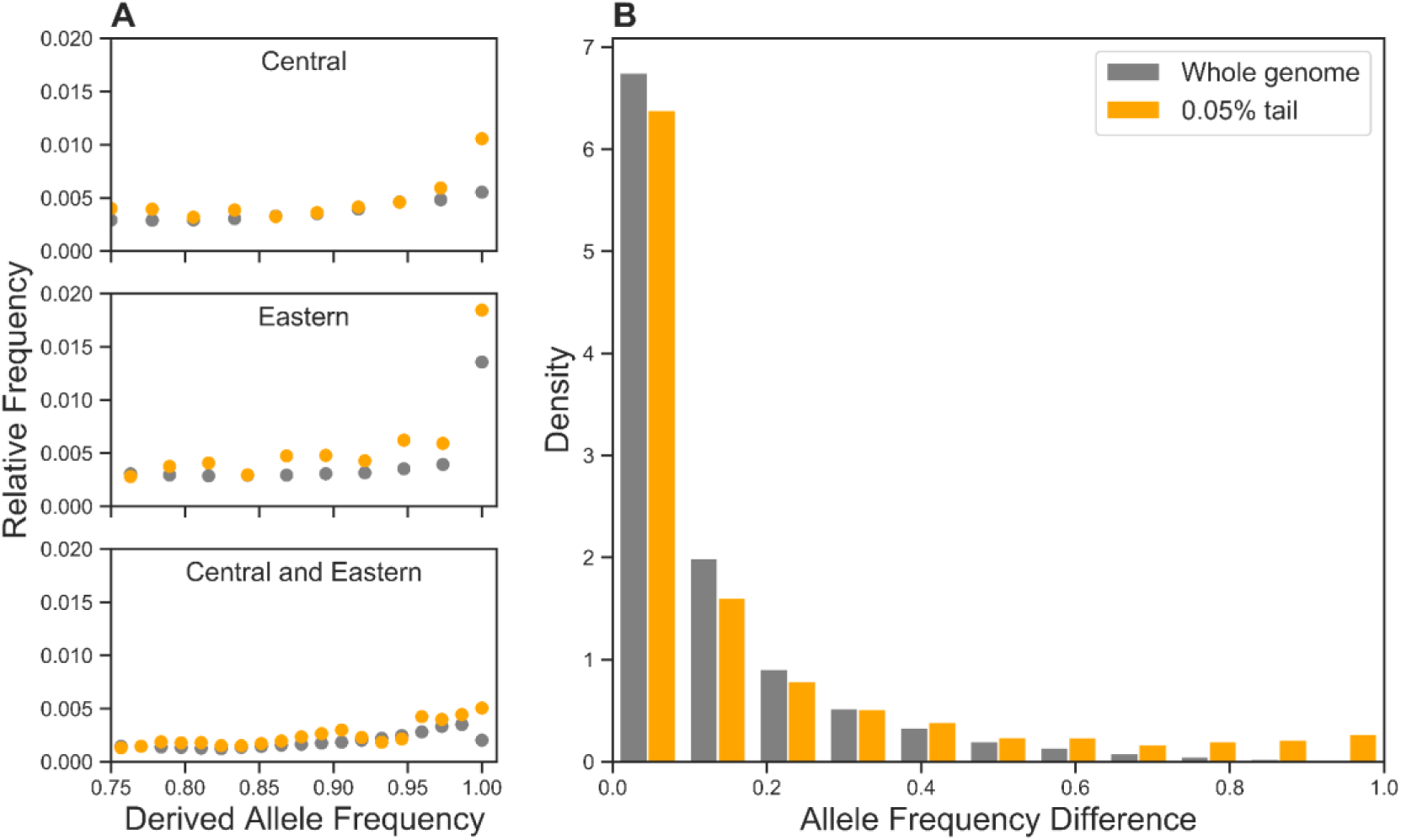
Site frequency patterns of SNPs in candidate windows. Allele frequencies of SNPs genome-wide (grey) and at the most stringent 3P-CLR tail threshold (0.05% candidates for positive selection in the central-eastern ancestor, orange). **A**: Unfolded SFS for central, eastern and central and eastern combined. The X axis is limited to focus on high-frequency derived alleles, full SFS in Figure S2, SFS at all 3P-CLR tail thresholds in Figure S3. **B**: Absolute DAF difference between central-eastern and Nigeria-Cameroon.

### SIV-related selection in the central-eastern ancestor

Under the hypothesis that SIV was a strong driver of adaptive evolution in the central-eastern ancestor, genes with signatures of positive selection would fall in known SIV-related functions more often than expected by chance. We test this expectation with enrichment tests for existing SIV/HIV-related categories on the genes that overlap our candidate windows, using Gowinda (Kofler and Schlotterer 2012). Gowinda tests for overrepresentation of a given gene set in our candidate windows compared to the expectation under neutrality (Kofler and Schlotterer 2012) (see Methods).

First we investigate selection on the immune response to SIV, by testing for an enrichment among the candidates in SIV-response genes, which in natural hosts change expression after SIV infection (Jacquelin et al. 2014; Jacquelin et al. 2009) and whose concerted action is thought to control SIV infection in natural hosts (Jacquelin et al. 2014; Jacquelin et al. 2009). SIV-response genes are enriched in signatures of positive selection in eastern chimpanzees (Schmidt et al. 2019) and vervet monkeys (Svardal et al. 2017), suggesting that they mediate adaptation to SIV. We find that the strongest candidate targets of positive selection in the central-eastern ancestor are modestly enriched in SIV-response genes, (0.05% candidates threshold, 11.8 expected, 18 observed, p-value=0.043; Figure 3). This signature can be refined by exploring enrichment in the 33 modules of differentially expressed genes that co-express temporally during SIV infection, defined by Svardal *et al.,* (Svardal et al. 2017). Two SIV co-expression modules are significantly enriched among candidate genes (Figure 3), both of which are defined by an acute response to SIV infection six days post-infection and exhibit strong signatures of positive selection in vervet monkeys (Svardal et al. 2017). Six days post-infection corresponds to when SIV can be first detected and when the immune response is typically initiated in natural hosts (Svardal et al. 2017).

**Figure 3.**
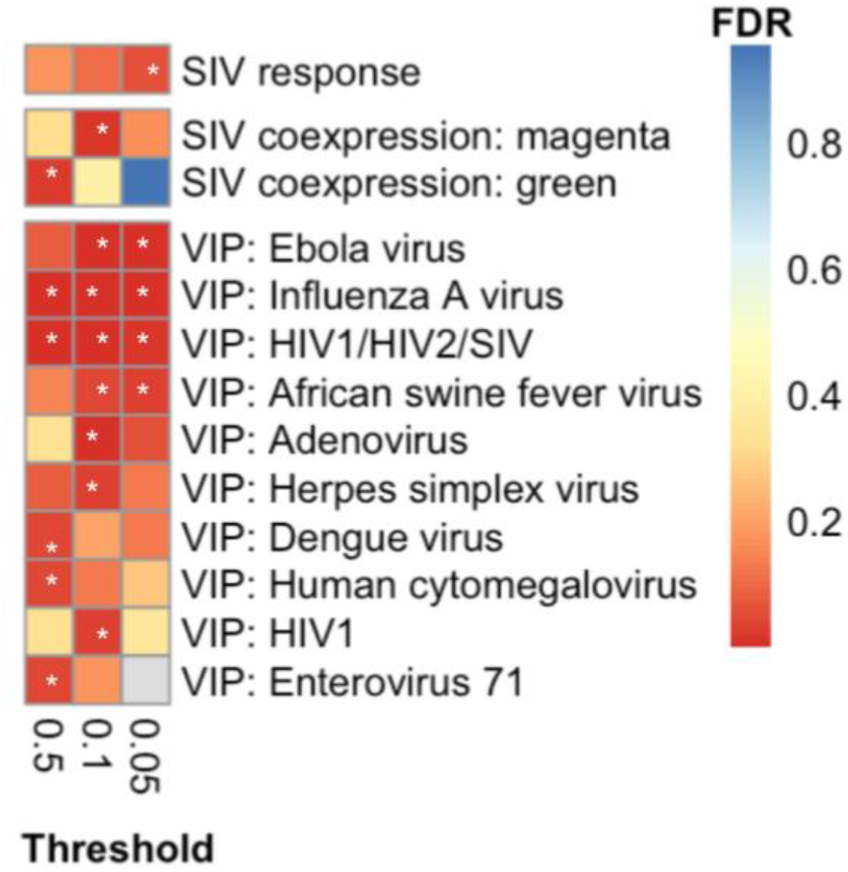
Enrichment of HIV/SIV-related and VIP categories in 3P-CLR candidate genes for the central-eastern ancestor. Columns for the three 3P-CLR quantiles, rows for each category that is significantly enriched in at least one quantile (FDR<0.05). From top to bottom: SIV response genes (Jacquelin et al. 2014, 2009), SIV co-expression modules (Svardal et al. 2017) and VIPs (Enard et al. 2016; Enard and Petrov 2018). For the SIV-response gene set there is only one category, hence we consider p-value<0.05 as significant, following (Schmidt et al. 2019). Colours represent FDR values, with grey representing categories undetected in that particular quantile. Stars mark categories with FDR < 0.05.

Next we investigate selection on host-virus physical protein interactions, by testing whether the candidates are enriched in genes that encode Viral Interacting Proteins (VIPs) –host proteins that physically interact with viral proteins, viral RNA or viral DNA (Enard et al. 2016; Enard and Petrov 2018). HIV/SIV-interacting VIPs have not been shown to be enriched in signatures of positive selection in chimpanzees or vervet monkeys, but VIPs are clear mediators of adaptation to viruses (Enard et al. 2016; Enard and Petrov 2018). Testing enrichment in all the 152 defined VIP categories, one of the strongest and most consistent enrichment signals is in VIPs that interact with HIV/SIV, across all candidate cut-offs (0.5% candidates threshold, p-value=0.00046; 0.1% candidates threshold, p-value=0.00014; 0.05% candidates threshold, p-value=0.00398; FDR values given in Figure 3). Interestingly, we find similarly strong enrichment in one additional category: influenza-interacting VIPs (0.5% candidates threshold, p-value=0.00002; 0.1% candidates threshold, p-value=0.00010; 0.05% candidates threshold, p-value=0.00014; FDR values given in Figure 3), in agreement with recent work in humans suggesting that RNA viruses are an important selective force in mammals (Enard et al. 2016). Given the differences in gene set sizes between the influenza and SIV/HIV (984 vs 810 genes) we consider the enrichment results largely similar. At the 0.05% tail for influenza we expect 4.568 genes and observe 14 (p-value = 0.00014, FDR=0.00112), whereas for HIV/SIV we expect 4.323 genes and observe 11 (p-value=0.00398, FDR=0.01854). Of note, these two categories overlap substantially, with 36% of the 0.05% tail candidate genes in the influenza VIP set being also HIV/SIV VIPs, which makes it difficult to establish the independence of their signatures. Not surprisingly other VIP categories also show some evidence of positive selection. Ebola is an interesting example, although it is a very small category with substantial overlap with HIV/SIV: three of the four 0.05% tail candidate genes in the ebola VIPs set are also HIV/SIV VIPs. Thus, evidence for other VIP categories exist, but together our results point to HIV/SIV as a particularly important selective force in chimpanzees.

Finally, we explore other biological categories in a hypothesis-free analysis. An enrichment test of GO categories (Ashburner et al. 2000) reveals 25 significantly enriched GO categories, two of which are relevant to host-viral interactions: ‘IκB/NFκB complex’ (0.5% candidates threshold, p-value=0.00008) and ‘positive regulation by host of viral transcription’ (0.5% candidates threshold, p-value=0.00002) (see rest of categories and FDR values in Figure S4). Notably, as discussed below, these two categories are intimately involved in the host biology under SIV/HIV infection. As expected, this analysis indicates likely adaptations to selective forces beyond SIV, although no category is as consistently enriched across thresholds as the HIV/SIV VIPs or SIV-response categories.

Thus, the potential targets of positive selection in the central-eastern ancestral population show enrichment patterns expected under adaptation to SIV/HIV and point to physical protein interactions (VIPs) and initial immune reaction (SIV-response) as likely being key in early adaptations to zoonotic SIV. Interestingly, the candidate windows are not enriched in existing HIV-related GWAS hits when enrichment is investigated with Gowinda as above. This suggests that while association studies of AIDS-related traits can identify genetic variants involved in clinical phenotypes (often with treatment), they do not reveal the genetic bases of the biological mechanisms that may allow natural hosts to control the pathogenicity of the virus.

### Potential sweeps of non-synonymous variants in *CD4*

A fundamental gene in SIV pathology is *CD4*, which encodes the glycoprotein required for HIV/SIV cell entry, additional to a chemokine coreceptor (Blumenthal et al. 2012). Excitingly, *CD4* lies within a candidate window at the 0.5% 3P-CLR threshold, showing signatures of positive selection in the central-eastern ancestral population. *CD4*’s protein-coding genomic region also contains SNPs with high derived allele frequencies in the central-eastern clade that are also highly differentiated compared with Nigeria-Cameroon (Figure 4A), as well as multiple signals of positive selection both in central and eastern chimpanzees (with PBSnj, data from Schmidt *et al.,* (Schmidt et al. 2019)) (Figure 4A). These signatures provide support for positive selection driving the evolutionary history of the gene in chimpanzees.

**Figure 4.**
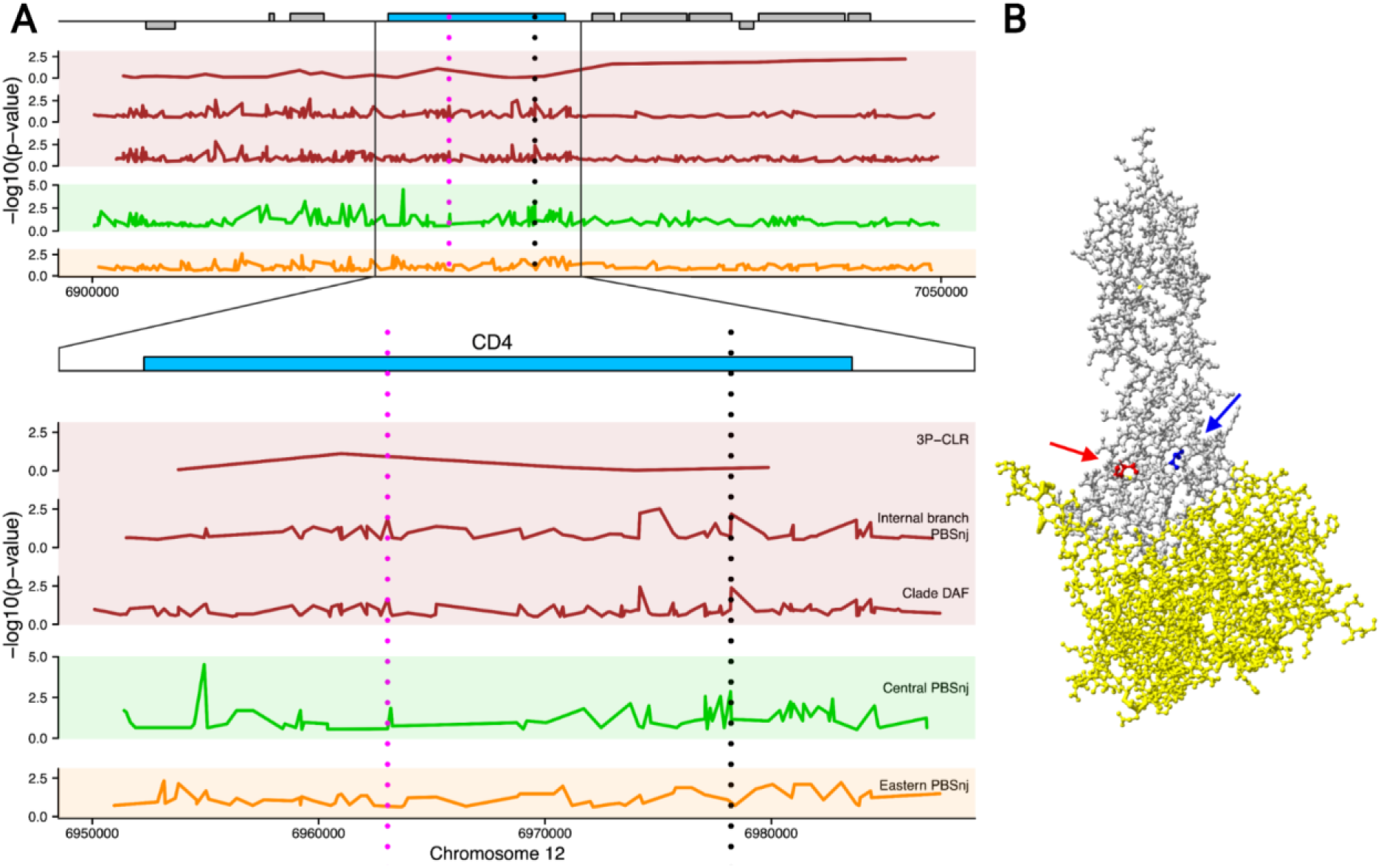
Signatures of selective sweeps in the extended *CD4* locus. **A.** Genomic representation of 150 kb and a magnified view of *CD4* with signatures of positive selection from 3P-CLR, PBSnj and clade-DAF. Statistics are coloured by population. The vertical dotted lines indicate best candidate SNPs of interest. The pink line indicates a splice site synonymous variant at position 6963043 with signatures of positive selection in the central-eastern ancestor. The black dotted line indicates two SNPs 39bp apart: a missense SNP in position 6978193 (the V55I SNP) with signatures of positive selection in centrals, and a missense SNP at position 6978232 (the P68T SNP) with signatures of selection in the central-eastern ancestor. For the DAF of these candidate SNPs across chimpanzee subspecies see Figure S5. **B.** Protein structure diagram showing the first two CD4 domains (grey) bound to the HIV envelope protein (yellow) (Kwong et al. 1998) and the amino acids corresponding to the two missense SNPs of interest, 6978193 (V55I) in blue and 6978232 (P68T) in red.

Bibollet-Ruche *et al.,* (Bibollet-Ruche et al. 2019) have shown that CD4 harbours functionally relevant variants in chimpanzees, so to integrate both types of information and better understand the gene’s signatures of selection we aimed to localise the most likely selected variant(s). We focused on SNPs with the highest allele frequency differentiation between the two chimpanzee clades (central-eastern and Nigeria-Cameroon-western), as they are prime candidates to explain the selective sweeps in the central-eastern ancestor identified by 3P-CLR (see Methods). We identified these SNPs with PBSnj in the internal branch (representing the central-eastern ancestor) with data from (Schmidt et al. 2019) and combine this with information on the amino acids found to determine SIV infectivity of chimpanzee cells by Bibollet-Ruche *et al.,* (Bibollet-Ruche et al. 2019). The P68T variant (chr12:6978232) falls in the tail of the empirical PBSnj distribution (PBSnj p-value=0.0066), and is among the SNPs with the highest PBSnj value in this window. It is one of only four CD4 amino acid polymorphisms known to determine SIV infectivity of chimpanzee cells (Bibollet-Ruche et al. 2019) being located, in the folded protein, near the binding site between CD4 and SIV’s *env* protein (Figure 4B). Central and eastern chimpanzees carry the ancestral P allele, able to inhibit SIVcpz. The P68T variant is a prime candidate to drive the selective sweep we observe in the central-eastern ancestor, with selection driving the P allele, though it remains possible that selection acted on a different genetic variant, or on multiple.

Interestingly, P68T is not the only *CD4* functional variant with signatures of selection in chimpanzees. The V55I variant (chr12:6978193) (Figure 4A) is not strongly differentiated in this branch (PBSnj p-value=0.1217), but has PBSnj signatures of positive selection in central chimpanzees (central PBSnj p-value = 0.0014). Like P68T, in the folded protein V55I is near the binding site between CD4 and SIV’s *env* protein (Figure 4B), and it is likely also functionally relevant, though the evidence from Bibollet-Ruche *et al.,* (Bibollet-Ruche et al. 2019) is weaker than for P68T. We note that in addition, a splice site region synonymous variant (chr12:6963043) is also among the most highly differentiated SNPs in the internal branch according to PBSnj (Figure 4B). It seems clear that CD4 being a key player in SIV infection, has been targeted by positive selection recurrently and in complex ways. Together with the three chemokine co-receptors showing signatures of positive selection in central chimpanzees (Schmidt et al. 2019), this indicates that proteins involved in SIV cell entry have evolved under adaptive evolution across chimpanzee populations.

### SIV-mediated selection across populations

Our results thus indicate that SIV was a strong selective force in the common ancestor of central and eastern chimpanzees. This adds to previous evidence of SIV-related adaptation both in the central and eastern populations (Schmidt et al. 2019). To understand how chimpanzees have been able to adapt to SIV thus requires integrating information across these populations. We integrated selection scores across the three populations (3P-CLR for the central-eastern ancestral population and previously computed PBSnj for the central and eastern subspecies (Schmidt et al. 2019), Figure 1) and tested as above if any SIV-related biological categories have been targeted by positive selection across populations using Gowinda (Figure S7).

The VIPs HIV/SIV category shows nominal enrichment across all possible configurations of populations: the central-eastern ancestor and central subspecies (p-value=0.00894, BH-FDR=0.185535), the central-eastern ancestor and eastern subspecies (p-value=0.00662, BH-FDR=0.1855350), and the three populations together (central-eastern ancestor, central and eastern subspecies; p-value=0.0161, BH-FDR=0.2700533). None of these are significant after correcting FDR values for the number of populations tested using a Benjamini and Hochberg (BH) correction; however, Gowinda is underpowered in this scenario, as several genes show signatures of positive selection in multiple lineages, but are counted only once. To address this limitation, we developed a custom method, ‘*set_perm*’, which extends the functionality of Gowinda to test if the joint distribution of candidate loci from different populations reveals enrichment for particular biological functions (Methods, Supplementary Note). Focusing on the three populations together, *set_perm* reveals significant enrichment (after correcting FDR values for testing multiple populations using a BH correction) for the VIPs HIV/SIV category (105.37901 expected, 145 observed, p-value=0.00003, BH FDR=0.00159), SIV-response genes (276.44037 expected, 308 observed, p-value=0.01678, BH FDR=0.04604) and the green SIV co-expression module (8.4254 expected, 20 observed, p-value=0.00039, BH FDR=0.02241) (Figure S7). This green co-expression module is one of the two showing evidence of positive selection in the central-eastern ancestor, and as mentioned above is associated with an acute response to SIV (Svardal et al. 2017). The higher power of *set_perm* when genes have signatures in several populations means that additional categories also become significant, including VIPs for influenza (112.18832 expected, 145 observed, p-value=0.00061, BH FDR=0.01815) and VIPs for dengue viruses (9.40297 expected, 20 observed, p-value=0.00083, BH FDR=0.02742).

To explore more generally which biological processes have been targeted by positive selection, we also tested for enrichment of signatures of positive selection in KEGG pathways (Qiu 2013). The KEGG pathway database consists of manually drawn molecular interaction diagrams for a wide range of biological pathways (Qiu 2013). Using Gowinda, only one pathway exhibits nominal enrichment in the combined candidates from the three populations (central-eastern ancestor, central and eastern subspecies): “Th1/Th2 cell differentiation pathway” (10.6 expected, 22 observed, p-value=0.00056, BH-FDR=0.25552) (Figure S8 and Table S1). Underpinning this signal are 22 genes: five with signatures of positive selection in the central-eastern ancestor, eleven in eastern and nine in central chimpanzees, of which three genes are candidates both in eastern and central chimpanzees (Figure 5). This result is confirmed and strengthened using *set_perm*, where this pathway shows a strongly significant enrichment even after correcting for testing multiple populations: (“Th1/Th2 cell differentiation pathway”, 10.2 expected, 25 observed, p-value=0.00002, BH-FDR=0.013800) (Figure S9). The similar contribution of signatures of selection in the three populations shows that the Th1/Th2 pathway has likely repeatedly undergone selection through time, although the individual genes targeted have largely differed across the three populations. This pathway is intimately associated with SIV infection, being active only in CD4+ T cells, which are the only cells that SIV infects (since CD4 is required for cell entry). Moreover, SIV coordinates its replication with T cell activation, as differentiation of CD4+ T helper cells involves upregulation of the transcription factors required to transcribe HIV/SIV genes (Cullen 1991; Mingyan et al. 2009). By this process, HIV replication leads to pathogenesis via the destruction of CD4+ T cells and the development of AIDS (Okoye and Picker 2013). Finally, the Th1/Th2 cell differentiation pathway is critical to generate an efficient immune response against viruses such as HIV/SIV (see discussion). Of note, multiple genes with signatures of positive selection in this pathway encode for proteins that play critical roles in SIV/HIV infection (Figure 5). These include the transcription factors Nuclear Factor Kappa B (NFκB) and Nuclear Factor of Activated T-cells (NFAT) (see Discussion).

**Figure 5.**
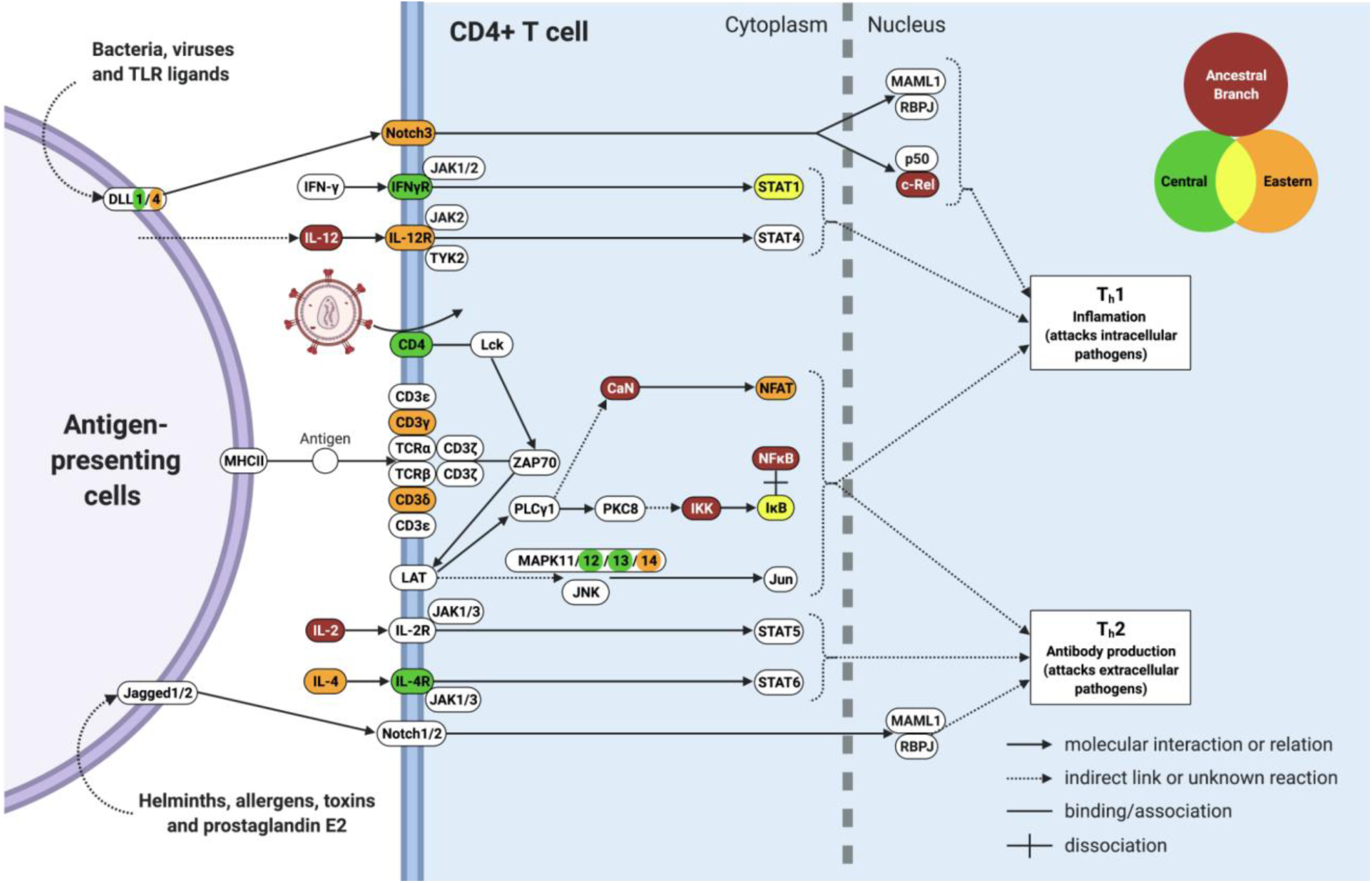
Simplified diagram of the Th1/Th2 cell differentiation pathway, highlighting proteins encoded by genes with signatures of positive selection. Proteins are coloured according to the population(s) in which they show signatures of positive selection (see legend), with white indicating proteins with no signatures of positive selection. Proteins in contact with other proteins indicate protein complexes. *IL13* is a candidate in both central and eastern but is not included in this simplified diagram. *RELA* is a candidate gene in the central-eastern ancestor and codes for a protein subunit of both NFκB and c-Rel. This diagram was created with BioRender.com.

The higher power of *set_perm* means that it reveals additional significantly enriched KEGG categories (Figure S9). For example, in the three populations together the immune-related ‘primary immunodeficiency’ category becomes significant (2.0645 expected, 10 observed, p-value=0.00003, BH FDR=0.006900), although it has only eight significant genes, half of them in centrals. Interestingly ‘Epstein-Barr virus infection’ (18.09451 expected, 37 observed, p-value=0.00003, BH FDR=0.013800) and ‘leishmaniasis’ (5.61591 expected, 17 observed, p-value=0.00006, BH FDR=0.016848) pathways also show significant enrichment in the combined three populations. These are not biological networks but rather pathogen-associated categories that might possibly point to additional selective pressures. Still, we note that 69.2% of the significant ‘leishmaniasis’ genes belong also to the larger Th1/Th2 cell differentiation pathway. Thus, to what extent this signature is fully independent from those in the Th1/Th2 cell differentiation pathway is unclear.

In summary, our genome-wide analysis shows strong evidence of positive selection in the common ancestor of central and eastern chimpanzees for SIV-response genes, SIV co-expression modules, and HIV/SIV VIPs (together with VIPs for other viruses). Further, when combining candidates from the central-eastern ancestor with those of its daughter species, we find evidence of positive selection in each of these SIV-related categories, together with recurrent positive selection in immune-related pathways, notably the “Th1/Th2 cell differentiation pathway”.

## Discussion

SIV is a formidable selective force, which has seemingly driven genetic adaptation in chimpanzees since zoonotic infection until recent times. Here we identify adaptations to SIV in the central-eastern ancestor. Together with our previous work showing evidence of positive selection in multiple SIV-related genes in central and eastern chimpanzees, but not in western and Nigeria-Cameroon (Schmidt et al. 2019), this supports the notion that zoonotic transmission took place in the central-eastern ancestor and that SIV immediately became a strong and continuous selective pressure in infected populations.

In the central-eastern ancestral population, loci with the strongest evidence of positive selection in the genome are significantly enriched in two sets of genes that are intimately associated with SIV, although in substantially different ways: genes that change expression upon SIV infection (‘SIV-response genes’ and ‘SIV co-expression modules’) and genes that encode proteins which physically interact with HIV/SIV (‘VIPs’). We also find evidence of positive selection driving sequence evolution of CD4, a protein with a critical role in immunity which is necessary for SIV cell entry. This reveals how diverse the initial adaptation to SIV likely was, consisting of genetic changes in host proteins involved in the host response, physical interaction and cell entry of SIV.

Notably, only the VIPs show an enrichment in signatures of positive selection only in the ancestral population, with the other categories showing evidence of selection at later times too. It is easy to imagine why VIPs may be under strong selective pressure after first infection, as they physically interact with the virus, while other mechanisms of adaptation may become more relevant at later times.

Genes involved in the host response to SIV in natural hosts (SIV-response genes) show evidence of positive selection not only in the central-eastern ancestral population, but also in eastern chimpanzees (Schmidt et al. 2019). Combined, these observations suggest that the modulation of the host response through gene expression evolves immediately after zoonotic transmission, and continues over time. This idea agrees well with the fact that SIV-response genes evolve under positive selection also in vervet monkeys (Svardal et al. 2017), which are long-term SIV natural hosts. It is interesting that SIV-response genes were not identified experimentally in chimpanzees, but in vervet monkeys, (Svardal et al. 2017; Jacquelin et al. 2014; Jacquelin et al. 2009). Their apparent role in adaptation in both species is consistent with the idea that host tolerance to SIV may evolve via similar mechanisms across distantly related species.

Genes that encode for the few proteins required by SIV to enter the cell also show evidence of positive selection across populations. Bibollet-Ruche *et al.,* (Bibollet-Ruche et al. 2019; Russell et al. 2021) showed that CD4, which is the required primary receptor for SIV cell entry, harbours multiple functionally relevant genetic variants. Complementing this important work, we find evidence of positive selection in the central-eastern ancestor at *CD4* with the most likely selected variant, P68T, which we hypothesise underwent a soft sweep in the central-eastern ancestor, being among only four CD4 missense variants shown to modify SIV infectivity in chimpanzee cells (Bibollet-Ruche et al. 2019). Of note, with both alleles chimpanzee P68T inhibits SIVcpz compared with the human CD4, though the derived T allele is most inhibitory (Bibollet-Ruche et al. 2019). The T allele is fixed in western chimpanzees while Nigeria-Cameroon chimpanzees are polymorphic (0.55) and bonobos and humans harbour the P allele, which is ancestral (Russell et al. 2021; Bibollet-Ruche et al. 2019). It is thus likely that the site was polymorphic in the common ancestor of all chimpanzees, making selection on P68 in the central-eastern ancestor likely to have occurred on standing variation–although it remains possible that selection on a different variant is responsible for the signatures of selection in CD4. It is nevertheless interesting that the site is invariant in western chimpanzees. Bibollet-Ruche *et al.,* (Bibollet-Ruche et al. 2019) proposed that fixation of the T allele in western chimpanzees could be due to adaptation to an SIV-related retrovirus. This is an interesting hypothesis that we are able to test using a formal test of positive selection in western chimpanzees, the PBSnj statistic in the western branch from Schmidt *et al.,* (Schmidt et al. 2019). Under our conservative 0.5% critical value western chimpanzees show no significant evidence of positive selection at P68T (PBSnj western branch p-value=0.0218). Still, this SNP falls within the 5% tail of the empirical distribution, and under less stringent criteria would be considered to harbour signatures of positive selection. Thus, while we cannot discard the possibility that the lack of diversity at P68T in western chimpanzees is explained by random genetic drift, which is strong in this subspecies due to their small effective population size and bottlenecks (de Manuel et al. 2016), our results suggest that the fixation of the T allele in western chimpanzees could be due to positive selection. CD4 has a critical role in immunity and has been suggested to have some signatures of positive selection in species without historical infection by SIV (e.g. humans (Meyerson et al. 2014)). Given the absence of natural SIV infections in western chimpanzees it appears more likely that a different virus, perhaps a related retrovirus, may have driven selection in this subspecies. In any case, we note that the allele frequency change in P68T in western chimpanzees has no effect over the signatures of positive selection in *CD4* in the central-eastern ancestor (we used Nigeria-Cameroon as the outgroup).

Beyond P68T, *CD4* harbours signatures of positive selection in other SNPs and subspecies. As mentioned in the Results, the V55I variant, which due to its position in the protein is likely to have functional effects, has PBSnj signatures of positive selection in central chimpanzees. Further, a C to T SNP within *CD4* (chr12:6973383) has significant signatures of positive selection in Nigeria-Cameroon (PBSnj p-value = 0.002) (data from (Schmidt et al. 2019)). This adds to a picture of multiple sweeps, both ancestral and lineage-specific, in *CD4*. Although outside of the scope of this paper, an extensive analysis of this complex region that integrates functional and evolutionary information would be extremely interesting. Remarkably, the multiple signatures of positive selection in *CD4* add to existing evidence of adaptation in central chimpanzees in *CCR3*, *CCR9* and *CXCR6* (Schmidt et al. 2019), which mediate SIV/HIV cell entry (Elliott et al. 2015; Wetzel et al. 2017; Nedellec et al. 2009) together with the primary receptor CD4.

Together, our results reveal that combining information across time and populations is critical to understand chimpanzee adaptation to SIV. By integrating selection scores from the central-eastern ancestor with those from the central and eastern subspecies, we identify strong enrichment in the Th1/Th2 cell differentiation pathway. Twenty-two genes in the Th1/Th2 cell differentiation pathway have signatures of positive selection across time, three genes in two separate populations. Together these observations show that this molecular pathway has been repeatedly hit by positive selection over time. Why this pathway? First, the Th1/Th2 cell differentiation pathway is critical for immunity against intracellular pathogens, including viruses. Naïve CD4+ T cells recognise a MHC class II peptide, are activated and divide to give rise to clone effector CD4+ T cells specific for that antigen (Zhang et al. 2014). CD4+ T cells can differentiate into T helper type-1 (Th1), T helper type-2 (Th2), or other T helper types, each with distinct cytokine-secretion phenotypes, production of distinct interferons, and different downstream immune responses (Zhang et al. 2014). By shaping which type of helper cell a CD4+ T cell will differentiate into, this pathway has critical effects on immunity.

Second, the Th1/Th2 cell differentiation pathway is particularly relevant for HIV/SIV pathogenesis, especially for control of viral replication. This is illustrated by the fact that a unique subset of humans, ‘HIV controllers’, who are able to spontaneously control HIV infection without treatment, are characterised by a Th1 differentiation bias (Vingert et al. 2012; Gomes et al. 2017). Specifically, HIV controllers differentiate more naïve CD4+ T cells into Th1 than into other T helper subsets (Gomes et al. 2017; Vingert et al. 2012). *In vitro* studies demonstrate that Th1 cells are more resistant to HIV replication than Th2 cells, since their higher expression of *APOBEC3G* limits reverse transcription and integration of HIV virions in Th1 better than in Th2 cells (Vetter and D’Aquila 2009; Kervevan and Chakrabarti 2021). It thus seems natural that biasing the Th1/Th2 cell differentiation pathway towards Th1 differentiation may be an efficient route to control SIV/HIV pathogenicity.

Identified candidate targets of selection include key transcription factors required for SIV/HIV replication: NFκB (in central-eastern ancestor) and NFAT (in easterns). NFAT proteins and HIV-1 upregulate each other and may allow establishment of HIV-1 in the early stages of infection (Pessler and Cron 2004). Both NFκB and NFAT bind to the same site which is identical in HIV-1 and HIV-2. We find signatures of positive selection in genes encoding proteins which inhibit (IκB in centrals and easterns) and activate (IKK in central-eastern ancestor) the NFκB protein complex. SIV/HIV have evolved to manipulate NFκB and IκB to minimise antiviral gene expression, while allowing NFκB-induced viral transcription (Hotter et al. 2017; Langer et al. 2019; Sauter et al. 2015). Hence, genes encoding NFκB, IκB and their immediate interactors are plausible targets for selection (Hotter et al. 2017; Langer et al. 2019; Sauter et al. 2015).

Our results contribute to the growing literature indicating that SIV has been and continues to be a strong selection pressure in chimpanzee evolution. Previous studies have largely focused on possible SIV-mediated selection by characterising variation at the MHC. Evidence of an ancient selective sweep at MHC-I in the ancestor of chimpanzees and bonobos was initially suggested to have been driven by SIV (de Groot et al. 2000; de Groot et al. 2002; de Groot et al. 2008) but is now attributed to an ‘SIV-like retrovirus’, given such an ancient SIVcpz origin (and subsequent loss of the virus in three lineages) is very unlikely (de Groot et al. 2017; Wroblewski et al. 2019). Whereas, evidence that SIV drives adaptation at MHC-I in the very recent past has been found by monitoring allele frequencies through time in eastern communities with high and low SIV loads (Wroblewski et al. 2015). We do not find evidence of positive selection at MHC-I in the central-eastern ancestor, although we likely have low power to identify signatures of selection in this highly complex region. Further, it is likely that balancing selection is acting on the MHC, as diversity at these sites can be protective against SIV and other viruses (Maibach et al. 2019), further complicating the picture. We note that a key novelty of our study is that the genomic dataset allowed us to go beyond the few genes known to play important roles in SIV/HIV infection and perform formal tests for selection genome-wide. Our approach provides compelling evidence of SIV-mediated selection only in the central-eastern chimpanzee clade.

Although we have focused on selection in response to SIV, additional selective pressures were surely relevant for the fitness of the central-eastern ancestor, and later populations. Here, in the central-eastern ancestor, we also identify significant enrichments of signatures of selection in VIPs that interact with influenza and other viruses, including ebolaviruses (Figure 3). Ebolaviruses have been implicated in disease and mortality of wild chimpanzees (Formenty et al. 1999). Influenza can infect captive chimpanzees (Buitendijk et al. 2014; Snyder et al. 1986; Kalter et al. 1997) and although respiratory and ‘flu-like’ diseases have been detected in wild chimpanzees, we are not aware of any study that has specifically detected influenza in such cases (Negrey et al. 2019; Hanamura et al. 2008; Kaur et al. 2008) and those would in any case represent contemporary infections. Nevertheless, it is possible that an archaic influenza-like virus infected chimpanzees thousands of years ago, leaving its mark in the genome in the same way as an unknown archaic SARS-CoV-2-like virus has been proposed to leave a signature of genetic adaptation in Asian human populations (Souilmi et al. 2021). In the future it would be interesting to explore the biological consequences of other targets of selection identified in our candidate windows.

Still, we find overwhelming evidence that SIV has been a strong selective force in central and eastern chimpanzees and their common ancestor, consistent with the scenario of zoonotic transmission of SIV into this ancestral population. This includes initial adaptations in SIV-interacting proteins, combined with adaptations on SIV response genes and cell entry genes that continued into daughter populations, and with adaptations in the Th1/Th2 cell differentiation pathway over time. This suggests the development of natural host immunity to SIV likely requires adaptation in the host-virus interacting factors and the factors that mediate cell entry and the immune response to infection. All of this strongly implies that SIV mediated selection has and continues to be important in the evolution of central and eastern chimpanzees, and that we can identify these adaptations with population genetics.

## Methods

### Simulations and Power Analysis

For 3P-CLR power analysis, we performed forward-in-time simulations in SLiM v.3.2 (Haller et al. 2019; Haller and Messer 2019). Following the inferred *Pan* demographic model (de Manuel et al. 2016; Won and Hey 2004) we simulated regions of length 1.2Mb, under neutrality and with positive selection. We assumed a 25-year generation time (Langergraber et al. 2012), 0.96×10^-8^ recombination rate (de Manuel et al. 2016; Won and Hey 2004) and a Wright-Fisher model. We performed 1,000 neutral simulations and 1,000 simulations for both selection coefficients (*s* = 0.1, 0.05). Our selection simulations followed a hard sweep, conditional on fixation before the central-eastern subspecies split (106 kya). Therefore, we sampled only a subset of all possible evolutionary trajectories of the beneficial mutation. Following Racimo (Racimo 2016) every 10^th^ SNP was a focal SNP, around which a 0.25cM window was centred, sliding every 10 SNPs. 100 SNPs were randomly sampled per window and used to calculate the 3P-CLR statistic, to test each window for selection signatures. For computational efficiency we extended 3P-CLR to allow the method to analyse 5kb segments of each chromosome. Results for the original and extended source codes show the expected high correlation (Figure S10). For the distribution of 3P-CLR scores for simulated under neutrality and positive selection see Figure S11. Nigeria-Cameroon was used as an outgroup to the central and eastern target populations. We used Nigeria-Cameroon rather than western who exhibit higher levels of drift and lower levels of segregating polymorphism (de Manuel et al. 2016). Hence using western as the outgroup would reduce the number of sites we could investigate, as 3P-CLR only considers sites which are segregating in the outgroup (Racimo 2016). We also note that gene flow between Nigeria-Cameroon and central-eastern chimpanzees may mask some signatures of selection, but would not generate them. We produced ROC curves to visualise 3P-CLR’s sensitivity and specificity, using pROC (Robin et al. 2011) in R v.3.5.2 [R Core Team 2018].

### Identifying selection in the central-eastern ancestor

We analysed genomic data generated by De Manuel et al. (de Manuel et al. 2016), using the same 3P-CLR parameters from our power analysis. This data consists of 47 genomes sequenced to high coverage (mean 22.5-fold coverage per individual), which had been sampled from chimpanzees of known subspecies: 18 central, 19 eastern, 10 Nigeria-Cameroon chimpanzees. We used the EPO alignment to infer the ancestral allele for calculating derived allele frequencies. We used the *Pan* diversity recombination map (Auton et al. 2012) to obtain genetic distances from physical distances. For the empirical distribution of 3P-CLR scores genome-wide and in the tails of the distribution see Figures S12-S13.

### *Post-hoc* SFS analysis of candidate windows in the central-eastern ancestor

We used the VCF file from De Manuel et al. (de Manuel et al. 2016) to calculate the unfolded SFS for the whole genome and for each 3P-CLR candidate window threshold. Sites were included only if they were polymorphic when considering central, eastern and Nigeria-Cameroon i.e. western specific variants were excluded. Combined central and eastern allele frequencies were calculated by simply pooling the samples as each subspecies had almost identical sample sizes (central: 18, eastern: 19). The difference in allele frequencies between central-eastern and Nigeria-Cameroon was calculated as the absolute difference in derived allele frequencies (DAF) between central-eastern and Nigeria-Cameroon for each site. The minimum DAF at which we can identify a polymorphism depends on the sample size. We have 18 central and 19 eastern samples, and so the minimum DAFs possible are as follows - central: 0.028, eastern: 0.026, central-eastern combined: 0.014.

### Gene set enrichment analyses in the central-eastern ancestor

We tested our 3P-CLR candidate windows for gene set enrichments, using Gowinda v.1.12 (Kofler and Schlötterer 2012), at the 3 highest quantiles of likelihood scores (0.5%, 0.1% and 0.05% candidate windows), as power to detect significant gene enrichment varies with the number of genes sampled. We tested for enrichment in different gene sets: SIV-response genes, SIV co-expression modules, VIPs, GO categories and KEGG biological pathways (Svardal et al. 2017; Enard et al. 2016; Enard and Petrov 2018; Ashburner et al. 2000; Jacquelin et al. 2014; Jacquelin et al. 2009; Qiu 2013). We note that the SIV co-expression modules are named as colours, consistent with (Svardal et al. 2017). Gowinda provides a p-value for the likelihood of finding the same evidence of selection seen in the candidate genes in a random set of genes, when considering genes of equivalent length and localisation in the genome (Kofler and Schlötterer 2012). These sets of random genes provide the appropriate negative controls for the analysis. Gowinda also calculates an FDR for each p-value. All genes tested were 1-1 human homologs and the analysis was run in ‘gene’ mode.

### Overlap with all PBSnj SNPs in the internal branch

To identify the SNPs with the strongest evidence of selection, we selected those with the largest allele frequency difference between the central-eastern and the western-Nigeria-Cameroon clades, within each significant 3P-CLR window at the 0.5% threshold. PBSnj (Schmidt et al. 2019) was used to assess allele frequency difference between the two clades. We then filtered by the 20 SNPs with the highest PBSnj value per window. This results in 10,676,926 SNPs genome-wide, which we annotated using ENSEMBL for pantro2.1.4, variant effect predictor (McLaren et al. 2016) and regulomeDB (Boyle et al. 2012). Functional annotation of the 20 highest PBSnj SNPs per window at the 0.5% threshold are shown in Figures S14-S15. SNPs of interest, additional to those identified in *CD4,* are shown in Table S2. We also calculated the difference in derived allele frequencies between the two clades (clade-DAF), as the difference in mean-weighted derived allele frequencies. The positions of amino acids within the structure of CD4 coded for by SNPs of interest were plotted using iCn3D v3.1.1 structure viewer (Wang et al. 2020).

### Identifying repeated targets of selection across branches

To investigate the targets of selection across time and populations, we combined selection candidates identified in the central-eastern ancestor with those previously identified in the central and the eastern subspecies by (Schmidt et al. 2019). To ensure each population is equally represented in the analysis, we assigned a single empirical p-value to each gene for each population and used genes in the 0.5% threshold for the central-eastern ancestor, to have an equal number of candidate genes as for the subspecies. Total number of candidate genes per population is as follows: in the central-eastern ancestor (817 genes), central p < 0.000192 (816 genes), eastern p < 0.000228 (806 genes). We tested for enrichment of our combined candidate sets in the same gene sets as above: SIV-response genes, SIV co-expression modules, VIPs, GO categories and KEGG biological pathways (Qiu 2013; Svardal et al. 2017; Jacquelin et al. 2014; Jacquelin et al. 2009; Enard et al. 2016; Enard and Petrov 2018; Ashburner et al. 2000) using Gowinda (Kofler and Schlötterer 2012). However, Gowinda is underpowered when testing for enrichment in multiple lineages.

In order to perform gene set enrichment tests on more than one lineage, we used a new method developed by JMS. ***Set_perm*** implements a permutation-based enrichment test that for a single lineage is equivalent to Gowinda (Kofler and Schlotterer 2012) but that correctly accounts for genes showing signatures of natural selection in more than one lineage. Details of the method can be found in the Supplementary Note and the code is available at https://github.com/joshuamschmidt/set_perm).

Briefly, a joint test of lineages is performed by summing the number of candidate genes per gene set across lineages. Thus, the total observed number of genes is simply the sum of the observed number of genes across the different lineages, per gene set. Note, that this means that the same gene can be counted more than once, if it is a candidate in more than one lineage.

Permutations are performed by first generating independent random gene sets for each lineage, and then combining them into a joint set across all lineages. Thus, In this case, for each gene set (S) and permutation (i), the joint permutation set, *joint_Si_*, is obtained by summing the number of genes (n) per lineage (k) i.e.

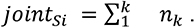

The p-values and FDR-corrected significance are calculated from permutation sets as described for Gowinda (Kofler and Schlotterer 2012).

As above, any gene present in more than one independent permutation test will contribute more than once to the joint permutation test.

Note, the number of observed genes may differ slightly between Gowinda and *set_perm*. This is because, in order to perform joint lineage testing with Gowinda we define candidates at the gene level, whereas with *set_perm* candidates are defined at the SNP level.

## Supporting Information

### Supplementary Files

**set_perm_SOM.docx: Supplementary note of *set_perm* method**

**enrichment_tables.xlsx: Supplementary tables of gene set enrichment results**

**SOM_RevisedMS_Pawar_ChimpSIV.docx: Supplementary figures S1-S15 and tables S1, S2**

### Supplementary Figures

**Figure S1: Power of 3P-CLR in the ancestral central-eastern population**. Each ROC curve was generated from 1000 neutral simulations and 1000 selection simulations for each s (s=0.05, 0.1).

**Figure S2: Full unfolded SFS for the whole genome and each 3P-CLR tail threshold in centrals (top left), easterns (top right) and central-eastern combined (bottom).** The central-eastern combined SFS was made by simply pooling all the samples as both subspecies have nearly identical sample sizes (central: 18, eastern: 19). The SFS are all indicative of selective sweeps.

**Figure S3. Site frequency spectrum of SNPs in candidate windows.** Allele frequencies of SNPs genome-wide and at different 3P-CLR tail thresholds. A: Unfolded SFS for central, eastern and central and eastern combined. The X axis is limited to focus on high-frequency derived alleles B: Absolute DAF difference between central-eastern and Nigeria-Cameroon.

**Figure S4: Enrichment of gene ontology (GO) categories across candidate genes at different 3P-CLR quantiles in the central-eastern ancestor**. Only categories with a significant enrichment in at least one quantile (FDR<0.05) are shown. Categories are separated by GO class: Biological Process, Cellular Component and Molecular Function. Colours represent FDR values (red as highest significance). Grey represents instances where a GO category was undetected in that particular quantile. Stars indicate a significant enrichment in that 3P-CLR quantile.

**Figure S5: DAF of the three candidate SNPs of interest in *CD4* across chimpanzee subspecies.** These candidate SNPs correspond to those highlighted in Figure 4. SNP at chr12:6963043 represents a splice variant with signatures of positive selection in the central-eastern ancestor. SNPs at chr12:6978193 (V55I SNP) and chr12:6978232 (P68T SNP) are missense variants with signatures of positive selection in centrals and the central-eastern ancestor respectively.

**Figure S6: Enrichment in SIV-related, VIP and GO categories of candidate targets of positive selection across populations, tested using Gowinda**. Abbreviations indicate the populations tested: central-eastern ancestor + central (A+C), central-eastern ancestor + eastern (A+E), central-eastern ancestor + central + eastern (A+C+E). Categories are separated by gene set tested: SIV responsive genes, SIV co-expression modules, VIPs and GO categories (Svardal et al. 2017; Enard et al. 2016; Enard and Petrov 2018; Ashburner et al. 2000; Jacquelin et al. 2014, 2009). Colours represent Benjamini and Hochberg corrected-FDR values (red as highest significance). We note that no category reaches significant enrichment (after correcting FDR values for the number of populations tested using a BH correction).

**Figure S7**: **Enrichment in SIV-related, and VIP categories of candidate targets of positive selection across populations, tested using *set_perm*.** Abbreviations indicate the populations tested: central (C), eastern (E), central-eastern ancestor (A), central-eastern ancestor + central (A+C), central-eastern ancestor + eastern (A+E), central-eastern ancestor + central + eastern (A+C+E). The following categories are significantly enriched in the three populations together at BH-corrected FDR < 0.05: SIV responsive genes, SIV co-expression green module, HIV/SIV VIPs, influenza (IAV) VIPs and dengue (DENV) VIPs. All of these categories are also significantly enriched when combining the central-eastern ancestor and the eastern subspecies. The SIV co-expression green module is significantly enriched when combining the central-eastern ancestor and the central subspecies. In the single lineages, we see significant enrichment in the eastern subspecies in the SIV-responsive and HIV/SIV VIPs . While the central-eastern ancestor is significantly enriched in the SIV co-expression green module, HIV/SIV VIPs and influenza (IAV) VIPs .

**Figure S8: Enrichment in KEGG pathways of candidate targets of positive selection across populations, tested using Gowinda**. Abbreviations indicate the population(s) tested: central-eastern ancestor (A), central-eastern ancestor + central + eastern (A+C+E). For the central-eastern ancestor we used candidate genes in the least stringent quantile (0.5), to match the number of candidates for the subspecies. Colours represent BH-corrected FDR values (red as highest significance). We note that none of the KEGG pathways reaches BH-corrected FDR < 0.05. The strongest enrichment is for the Th1 and Th2 cell differentiation pathway for A+C+E with p-value=0.00056 and BH-FDR=0.25552, which represents a nominal enrichment.

**Figure S9: Enrichment in KEGG pathways of candidate targets of positive selection across populations, tested using *set_perm*.** Abbreviations indicate the populations tested: central-eastern ancestor + central (A+C), central-eastern ancestor + eastern (A+E), central-eastern ancestor + central + eastern (A+C+E). The following KEGG categories are significantly enriched in the three populations together at BH-corrected FDR < 0.05: ‘Th1 and Th2 cell differentiation’, ‘primary immunodeficiency’, ‘Epstein-Barr virus infection’ and ‘leishmaniasis’. The ‘primary immunodeficiency’ and ‘leishmaniasis’ categories are also significantly enriched in the central subspecies. Hence for the ‘primary immunodeficiency’ and ‘leishmaniasis’ gene sets, the central lineage is likely driving the signal when the three populations are combined. When the central-eastern ancestor and eastern subspecies are combined, we see significant enrichment in the ‘chromosome and associated proteins’ pathway, which is also significantly enriched in the central-eastern ancestor.

**Figure S10: Correlation between the 3P-CLR values with the original and extended code.** Likelihood scores are significantly correlated between the original and modified 3P-CLR source code (rho=0.4565, p<2.2e-16). The absence of a perfect correlation is not due to differences in the algorithm, but due to the sampling variance of SNPs in each window. Specifically, if more than 100 SNPs are present within a given window, 3P-CLR chooses 100 SNPs at random. Thus, the same SNPs will not be sampled each time the method is run. Variation in 3P-CLR likelihood ratio scores therefore results if different SNPs were used to calculate the statistic. As expected, the correlation is weaker for low 3P-CLR but high at higher 3P-CLR values, where candidates of positive selection fall.

**Figure S11: Distribution of 3P-CLR scores** for data simulated under neutrality (A) and positive selection with selection coefficients of 0.05 (B) and 0.1 (C), 1000 replicates were generated in each case.

**Figure S12: Empirical distribution of 3P-CLR scores**. Vertical lines indicate the thresholds used to define candidate windows (0.5%, 0.1% and 0.05% tails of the empirical distribution).

**Figure S13: Distribution of 3P-CLR scores in the tails of the empirical distribution.** (A) the 0.5%, (B) 0.1% and (C) 0.05% candidate quantiles, corresponding to 4090, 818 and 409 genomic windows respectively.

**Figure S14: Functional annotation of 20 highest PBSnj SNPs per window at the 0.5% threshold using VEP.**

**Figure S15: Functional annotation of 20 highest PBSnj SNPs per window at the 0.5% threshold using regulomeDB**.

### Supplementary Tables

**Table S1: Number of candidate genes from each population which belong to gene categories of interest.** The number of genes which are candidates in central only, eastern only, and ancestral only are shown in the C, E and A columns respectively. The number of genes which are candidates in both central and eastern, central and ancestral, eastern and ancestral, or all three populations are shown in the C+E, C+A, E+A, and C+E+A columns respectively.

**Table S2: SNPs of interest in genes identified within candidate windows of positive selection at the 0.5% threshold in the central-eastern ancestor, additional to *CD4.*** SNPs with regulomeDB scores of 1f, 2a, 2b, 2c are considered to have putative significant regulatory function, due to the lack of eQTL data for chimpanzees. Category indicates the gene set each gene belongs to. Abbreviations: HCMV and ADV indicate human cytomegalovirus and adenovirus respectively.

## Supporting information

Supplementary note of set_perm method

Supplementary tables of gene set enrichment results

## Acknowledgments

We thank Dean Cornish, Clàudia Fontserè, Sojung Han, Martin Kuhlwilm and Tomàs Marquès-Bonet for helpful comments on the manuscript. HP was supported by a Formació de Personal Investigador fellowship from Generalitat de Catalunya (FI_B100131), HO by a London NERC DTP studentship (NE/S007229/1), and JS and AA by a UCL’s Wellcome Trust ISSF3 award (Grant Reference 204841/Z/16/Z).

## References

1. Ashburner, Michael, Catherine A. Ball, Judith A. Blake, David Botstein, Heather Butler, J. Michael Cherry, Allan P. Davis, et al. 2000. “Gene Ontology: Tool for the Unification of Biology. The Gene Ontology Consortium.” Nature Genetics 25 (1): 25–29.

2. Auton, Adam, Adi Fledel-Alon, Susanne Pfeifer, Oliver Venn, Laure Ségurel, Teresa Street, Ellen M. Leffler, et al. 2012. “A Fine-Scale Chimpanzee Genetic Map from Population Sequencing.” Science 336 (6078): 193–98.

3. Bailes, Elizabeth, Feng Gao, Frederic Bibollet-Ruche, Valerie Courgnaud, Martine Peeters, Preston A. Marx, Beatrice H. Hahn, and Paul M. Sharp. 2003. “Hybrid Origin of SIV in Chimpanzees.” Science 300 (5626): 1713.

4. Barouch, Dan H., Khader Ghneim, William J. Bosche, Yuan Li, Brian Berkemeier, Michael Hull, Sanghamitra Bhattacharyya, et al. 2016. “Rapid Inflammasome Activation Following Mucosal SIV Infection of Rhesus Monkeys.” Cell 165 (3): 656–67.

5. Bell, Sidney M., and Trevor Bedford. 2017. “Modern-Day SIV Viral Diversity Generated by Extensive Recombination and Cross-Species Transmission.” PLoS Pathogens 13 (7): e1006466.

6. Berger, Edward A. 1997. “HIV Entry and Tropism: The Chemokine Receptor Connection.” AIDS 11 Suppl A: S3–16.

7. Bibollet-Ruche, Frederic, Ronnie M. Russell, Weimin Liu, Guillaume B. E. Stewart-Jones, Scott Sherrill-Mix, Yingying Li, Gerald H. Learn, et al. 2019. “CD4 Receptor Diversity in Chimpanzees Protects against SIV Infection.” Proceedings of the National Academy of Sciences of the United States of America 116 (8): 3229–38.

8. Blumenthal, Robert, Stewart Durell, and Mathias Viard. 2012. “HIV Entry and Envelope Glycoprotein-Mediated Fusion.” The Journal of Biological Chemistry 287 (49): 40841–49.

9. Boyle, Alan P., Eurie L. Hong, Manoj Hariharan, Yong Cheng, Marc A. Schaub, Maya Kasowski, Konrad J. Karczewski, et al. 2012. “Annotation of Functional Variation in Personal Genomes Using RegulomeDB.” Genome Research 22 (9): 1790–97.

10. Buitendijk, Hester, Zahra Fagrouch, Henk Niphuis, Willy M. Bogers, Kristin S. Warren, and Ernst J. Verschoor. 2014. “Retrospective Serology Study of Respiratory Virus Infections in Captive Great Apes.” Viruses 6 (3): 1442–53.

11. Chahroudi, Ann, Steven E. Bosinger, Thomas H. Vanderford, Mirko Paiardini, and Guido Silvestri. 2012. “Natural SIV Hosts: Showing AIDS the Door.” Science 335 (6073): 1188–93.

12. Chimpanzee Sequencing and Analysis Consortium. 2005. “Initial Sequence of the Chimpanzee Genome and Comparison with the Human Genome.” Nature 437 (7055): 69–87.

13. Cullen, Bryan R. 1991. “Regulation of HIV-1 Gene Expression.” The FASEB Journal 5 (10): 2361–68.

14. Elliott, Sarah T. C., Katherine S. Wetzel, Nicholas Francella, Steven Bryan, Dino C. Romero, Nadeene E. Riddick, Farida Shaheen, et al. 2015. “Dualtropic CXCR6/CCR5 Simian Immunodeficiency Virus (SIV) Infection of Sooty Mangabey Primary Lymphocytes: Distinct Coreceptor Use in Natural versus Pathogenic Hosts of SIV.” Journal of Virology 89 (18): 9252–61.

15. Enard, David, Le Cai, Carina Gwennap, and Dmitri A. Petrov. 2016. “Viruses Are a Dominant Driver of Protein Adaptation in Mammals.” eLife 5: e12469.

16. Enard, David, and Dmitri A. Petrov. 2018. “Evidence That RNA Viruses Drove Adaptive Introgression between Neanderthals and Modern Humans.” Cell 175 (2): 360–71.e13.

17. Etienne, Lucie, Eric Nerrienet, Matthew LeBreton, Godwin Tafon Bibila, Yacouba Foupouapouognigni, Dominique Rousset, Ahmadou Nana, et al. 2011. “Characterization of a New Simian Immunodeficiency Virus Strain in a Naturally Infected Pan Troglodytes Troglodytes Chimpanzee with AIDS Related Symptoms.” Retrovirology 8: 4.

18. Formenty, Pierre, Christophe Boesch, Monique Wyers, Claudia Steiner, Franca Donati, Frédéric Dind, Francine Walker, and Bernard Le Guenno. 1999. “Ebola Virus Outbreak among Wild Chimpanzees Living in a Rain Forest of Côte d’Ivoire.” The Journal of Infectious Diseases 179 Suppl 1: S120–26.

19. Gilden, Raymond V., Larry O. Arthur, W. Gerard Robey, John C. Kelliher, Charles E. Graham, and Peter J. Fischinger. 1986. “HTLV-III Antibody in a Breeding Chimpanzee Not Experimentally Exposed to the Virus.” The Lancet 1 (8482): 678–79.

20. Goldstein, Simoy, Ilnour Ourmanov, Charles R. Brown, Ronald Plishka, Alicia Buckler-White, Russell Byrum, and Vanessa M. Hirsch. 2005. “Plateau Levels of Viremia Correlate with the Degree of CD4+-T-Cell Loss in Simian Immunodeficiency Virus SIVagm-Infected Pigtailed Macaques: Variable Pathogenicity of Natural SIVagm Isolates.” Journal of Virology 79 (8): 5153–62.

21. Gomes, Samara Tatielle M., Érica R. Gomes, Mike B. Dos Santos, Sandra S. Lima, Maria Alice F. Queiroz, Luiz Fernando A. Machado, Izaura M. V. Cayres-Vallinoto, Antonio Carlos R. Vallinoto, Marluísa de O Guimarães Ishak, and Ricardo Ishak. 2017. “Immunological and Virological Characterization of HIV-1 Viremia Controllers in the North Region of Brazil.” BMC Infectious Diseases 17 (1): 381.

22. Greenwood, Edward J. D., Fabian Schmidt, Ivanela Kondova, Henk Niphuis, Vida L. Hodara, Leah Clissold, Kirsten McLay, et al. 2015. “Simian Immunodeficiency Virus Infection of Chimpanzees (Pan Troglodytes) Shares Features of Both Pathogenic and Non-Pathogenic Lentiviral Infections.” PLoS Pathogens 11 (9): e1005146.

23. Groot, Natasja G. de, Corinne M. C. Heijmans, and Ronald E. Bontrop. 2017. “AIDS in Chimpanzees: The Role of MHC Genes.” Immunogenetics 69 (8-9): 499–509.

24. Groot, Natasja G. de, Corrine M. C. Heijmans, Nanine de Groot, Nel Otting, Annemiek J. M. de Vos-Rouweller, Edmond J. Remarque, Maxime Bonhomme, Gaby G. M. Doxiadis, Brigitte Crouau-Roy, and Ronald E. Bontrop. 2008. “Pinpointing a Selective Sweep to the Chimpanzee MHC Class I Region by Comparative Genomics.” Molecular Ecology 17 (8): 2074–88.

25. Groot, Natasja G. de, Nel Otting, Gaby G. M. Doxiadis, Sunita S. Balla-Jhagjhoorsingh, Jonathan L. Heeney, Jon J. van Rood, Pascal Gagneux, and Ronald E. Bontrop. 2002. “Evidence for an Ancient Selective Sweep in the MHC Class I Gene Repertoire of Chimpanzees.” Proceedings of the National Academy of Sciences of the United States of America 99 (18): 11748–53.

26. Groot, Natasja G. de, Nel Otting, Rafael Argüello, David I. Watkins, Gaby G. Doxiadis, J. Alejandro Madrigal, and Ronald E. Bontrop. 2000. “Major Histocompatibility Complex Class I Diversity in a West African Chimpanzee Population: Implications for HIV Research.” Immunogenetics 51 (6): 398–409.

27. Haller, Benjamin C., Jared Galloway, Jerome Kelleher, Philipp W. Messer, and Peter L. Ralph. 2019. “Tree-Sequence Recording in SLiM Opens New Horizons for Forward-Time Simulation of Whole Genomes.” Molecular Ecology Resources 19 (2): 552–66.

28. Haller, Benjamin C., and Philipp W. Messer. 2019. “SLiM 3: Forward Genetic Simulations Beyond the Wright-Fisher Model.” Molecular Biology and Evolution 36 (3): 632–37.

29. Hanamura, Shunkichi, Mieko Kiyono, Magdalena Lukasik-Braum, Titus Mlengeya, Mariko Fujimoto, Michio Nakamura, and Toshisada Nishida. 2008. “Chimpanzee Deaths at Mahale Caused by a Flu-like Disease.” Primates; Journal of Primatology 49 (1): 77–80.

30. Hotter, Dominik, Teresa Krabbe, Elisabeth Reith, Ali Gawanbacht, Nadia Rahm, Ahidjo Ayouba, Benoît Van Driessche, et al. 2017. “Primate Lentiviruses Use at Least Three Alternative Strategies to Suppress NF-κB-Mediated Immune Activation.” PLoS Pathogens 13 (8): e1006598.

31. Jacquelin, Béatrice, Véronique Mayau, Brice Targat, Anne-Sophie Liovat, Désirée Kunkel, Gaël Petitjean, Marie-Agnès Dillies, et al. 2009. “Nonpathogenic SIV Infection of African Green Monkeys Induces a Strong but Rapidly Controlled Type I IFN Response.” Journal of Clinical Investigation 119 (12): 3544–55.

32. Jacquelin, Béatrice, Gaël Petitjean, Désirée Kunkel, Anne-Sophie Liovat, Simon P. Jochems, Kenneth A. Rogers, Mickaël J. Ploquin, et al. 2014. “Innate Immune Responses and Rapid Control of Inflammation in African Green Monkeys Treated or Not with Interferon-Alpha during Primary SIVagm Infection.” PLoS Pathogens 10 (7): e1004241.

33. Kalter, Seymour S., Richard L. Heberling, Anthony W. Cooke, John D. Barry, Pei Y. Tian, and William J. Northam. 1997. “Viral Infections of Nonhuman Primates.” Laboratory Animal Science 47 (5): 461–67.

34. Kaur, Taranjit, Jatinder Singh, Suxiang Tong, Charles Humphrey, Donna Clevenger, Wendy Tan, Brian Szekely, et al. 2008. “Descriptive Epidemiology of Fatal Respiratory Outbreaks and Detection of a Human-Related Metapneumovirus in Wild Chimpanzees (Pan Troglodytes) at Mahale Mountains National Park, Western Tanzania.” American Journal of Primatology 70 (8): 755–65.

35. Keele, Brandon F., James Holland Jones, Karen A. Terio, Jacob D. Estes, Rebecca S. Rudicell, Michael L. Wilson, Yingying Li, et al. 2009. “Increased Mortality and AIDS-like Immunopathology in Wild Chimpanzees Infected with SIVcpz.” Nature 460 (7254): 515–19.

36. Keele, Brandon F., Fran Van Heuverswyn, Yingying Li, Elizabeth Bailes, Jun Takehisa, Mario L. Santiago, Frederic Bibollet-Ruche, et al. 2006. “Chimpanzee Reservoirs of Pandemic and Nonpandemic HIV-1.” Science 313 (5786): 523–26.

37. Kervevan, Jérôme, and Lisa A. Chakrabarti. 2021. “Role of CD4+ T Cells in the Control of Viral Infections: Recent Advances and Open Questions.” International Journal of Molecular Sciences 22 (2): 523.

38. Kofler, Robert, and Christian Schlötterer. 2012. “Gowinda: Unbiased Analysis of Gene Set Enrichment for Genome-Wide Association Studies.” Bioinformatics 28 (15): 2084–85.

39. Kwong, Peter D., Richard Wyatt, James Robinson, Raymond W. Sweet, Joseph Sodroski, and Wayne A. Hendrickson. 1998. “Structure of an HIV gp120 Envelope Glycoprotein in Complex with the CD4 Receptor and a Neutralizing Human Antibody.” Nature 393 (6686): 648–59.

40. Langergraber, Kevin E., Kay Prüfer, Carolyn Rowney, Christophe Boesch, Catherine Crockford, Katie Fawcett, Eiji Inoue, et al. 2012. “Generation Times in Wild Chimpanzees and Gorillas Suggest Earlier Divergence Times in Great Ape and Human Evolution.” Proceedings of the National Academy of Sciences of the United States of America 109 (39): 15716–21.

41. Langer, Simon, Christian Hammer, Kristina Hopfensperger, Lukas Klein, Dominik Hotter, Paul D. De Jesus, Kristina M. Herbert, et al. 2019. “HIV-1 Vpu Is a Potent Transcriptional Suppressor of NF-κB-Elicited Antiviral Immune Responses.” eLife 8: e41930.

42. Leitner, Thomas, Marie-Christine Dazza, Michel Ekwalanga, Cristian Apetrei, and Sentob Saragosti. 2007. “Sequence Diversity among Chimpanzee Simian Immunodeficiency Viruses (SIVcpz) Suggests That SIVcpzPts Was Derived from SIVcpzPtt through Additional Recombination Events.” AIDS Research and Human Retroviruses 23 (9): 1114–18.

43. Locatelli, Sabrina, Ryan J. Harrigan, Paul R. Sesink Clee, Matthew W. Mitchell, Kurt A. McKean, Thomas B. Smith, and Mary Katherine Gonder. 2016. “Why Are Nigeria-Cameroon Chimpanzees (Pan Troglodytes Ellioti) Free of SIVcpz Infection?” PloS One 11 (8): e0160788.

44. MacFie, Tammie S., Eric Nerrienet, Natasja G. de Groot, Ronald E. Bontrop, and Nicholas I. Mundy. 2009. “Patterns of Diversity in HIV-Related Loci among Subspecies of Chimpanzee: Concordance at CCR5 and Differences at CXCR4 and CX3CR1.” Molecular Biology and Evolution 26 (4): 719–27.

45. Ma, Dongzhu, Anna J. Jasinska, Felix Feyertag, Viskam Wijewardana, Jan Kristoff, Tianyu He, Kevin Raehtz, et al. 2014. “Factors Associated with Siman Immunodeficiency Virus Transmission in a Natural African Nonhuman Primate Host in the Wild.” Journal of Virology 88(10): 5687–705.

46. Ma, Dongzhu, Anna J. Jasinska, Jan Kristoff, J. Paul Grobler, Trudy Turner, Yoon Jung, Christopher Schmitt, et al. 2013. “SIVagm Infection in Wild African Green Monkeys from South Africa: Epidemiology, Natural History, and Evolutionary Considerations.” PLoS Pathogens 9 (1): e1003011.

47. Maibach, Vincent, Kevin Langergraber, Fabian H. Leendertz, Roman M. Wittig, and Linda Vigilant. 2019. “Differences in MHC-B Diversity and KIR Epitopes in Two Populations of Wild Chimpanzees.” Immunogenetics 71 (10): 617–33.

48. Mandell, Daniel T., Jan Kristoff, Thaidra Gaufin, Rajeev Gautam, Dongzhu Ma, Netanya Sandler, George Haret-Richter, et al. 2014. “Pathogenic Features Associated with Increased Virulence upon Simian Immunodeficiency Virus Cross-Species Transmission from Natural Hosts.” Journal of Virology 88 (12): 6778–92.

49. Manuel, Marc de, Martin Kuhlwilm, Peter Frandsen, Vitor C. Sousa, Tariq Desai, Javier Prado-Martinez, Jessica Hernandez-Rodriguez, et al. 2016. “Chimpanzee Genomic Diversity Reveals Ancient Admixture with Bonobos.” Science 354 (6311): 477–81.

50. McLaren, William, Laurent Gil, Sarah E. Hunt, Harpreet Singh Riat, Graham R. S. Ritchie, Anja Thormann, Paul Flicek, and Fiona Cunningham. 2016. “The Ensembl Variant Effect Predictor.” Genome Biology 17 (1): 122.

51. Meyerson, Nicholas R., Paul A. Rowley, Christina H. Swan, Dona T. Le, Gregory K. Wilkerson, and Sara L. Sawyer. 2014. “Positive Selection of Primate Genes That Promote HIV-1 Replication.” Virology 454–455 (April): 291–98.

52. Mingyan, Yu, Liu Xinyong, and Erik De Clercq. 2009. “NF-kappaB: The Inducible Factors of HIV-1 Transcription and Their Inhibitors.” Mini Reviews in Medicinal Chemistry 9 (1): 60–69.

53. Moore, John P., Scott G. Kitchen, Pavel Pugach, and Jerome A. Zack. 2004. “The CCR5 and CXCR4 Coreceptors—Central to Understanding the Transmission and Pathogenesis of Human Immunodeficiency Virus Type 1 Infection.” AIDS Research and Human Retroviruses 20 (1): 111–26.

54. Nedellec, Rebecca, Mia Coetzer, Naoki Shimizu, Hiroo Hoshino, Victoria R. Polonis, Lynn Morris, Ulrika E. A. Mårtensson, James Binley, Julie Overbaugh, and Donald E. Mosier. 2009. “Virus Entry via the Alternative Coreceptors CCR3 and FPRL1 Differs by Human Immunodeficiency Virus Type 1 Subtype.” Journal of Virology 83 (17): 8353–63.

55. Negrey, Jacob D., Rachna B. Reddy, Erik J. Scully, Sarah Phillips-Garcia, Leah A. Owens, Kevin E. Langergraber, John C. Mitani, et al. 2019. “Simultaneous Outbreaks of Respiratory Disease in Wild Chimpanzees Caused by Distinct Viruses of Human Origin.” Emerging Microbes & Infections 8 (1): 139–49.

56. Okoye, Afam A., and Louis J. Picker. 2013. “CD4(+) T-Cell Depletion in HIV Infection: Mechanisms of Immunological Failure.” Immunological Reviews 254 (1): 54–64.

57. Pessler, Frank, and Randy Q. Cron. 2004. “Reciprocal Regulation of the Nuclear Factor of Activated T Cells and HIV-1.” Genes and Immunity 5 (3): 158–67.

58. Prado-Martinez, Javier, Peter H. Sudmant, Jeffrey M. Kidd, Heng Li, Joanna L. Kelley, Belen Lorente-Galdos, Krishna R. Veeramah, et al. 2013. “Great Ape Genetic Diversity and Population History.” Nature 499 (7459): 471–75.

59. Prüfer, Kay, Cesare de Filippo, Steffi Grote, Fabrizio Mafessoni, Petra Korlević, Mateja Hajdinjak, Benjamin Vernot, et al. 2017. “A High-Coverage Neandertal Genome from Vindija Cave in Croatia.” Science 358 (6363): 655–58.

60. Qiu, Yu-Qing. 2013. “KEGG Pathway Database.” Encyclopedia of Systems Biology. Springer, New York, NY. https://doi.org/10.1007/978-1-4419-9863-7_472.

61. Racimo, Fernando. 2016. “Testing for Ancient Selection Using Cross-Population Allele Frequency Differentiation.” Genetics 202 (2): 733–50.

62. Robin, Xavier, Natacha Turck, Alexandre Hainard, Natalia Tiberti, Frédérique Lisacek, Jean-Charles Sanchez, and Markus Müller. 2011. “pROC: An Open-Source Package for R and S to Analyze and Compare ROC Curves.” BMC Bioinformatics 12: 77.

63. Rudicell, Rebecca S., James Holland Jones, Emily E. Wroblewski, Gerald H. Learn, Yingying Li, Joel D. Robertson, Elizabeth Greengrass, et al. 2010. “Impact of Simian Immunodeficiency Virus Infection on Chimpanzee Population Dynamics.” PLoS Pathogens 6 (9): e1001116.

64. Russell, Ronnie M., Frederic Bibollet-Ruche, Weimin Liu, Scott Sherrill-Mix, Yingying Li, Jesse Connell, Dorothy E. Loy, et al. 2021. “CD4 Receptor Diversity Represents an Ancient Protection Mechanism against Primate Lentiviruses.” Proceedings of the National Academy of Sciences of the United States of America 118 (13): e2025914118.

65. Santiago, Mario L., Cynthia M. Rodenburg, Shadrack Kamenya, Frederic Bibollet-Ruche, Feng Gao, Elizabeth Bailes, Sreelatha Meleth, et al. 2002. “SIVcpz in Wild Chimpanzees.” Science 295 (5554): 465.

66. Sauter, Daniel, Dominik Hotter, Benoît Van Driessche, Christina M. Stürzel, Silvia F. Kluge, Steffen Wildum, Hangxing Yu, et al. 2015. “Differential Regulation of NF-κB-Mediated Proviral and Antiviral Host Gene Expression by Primate Lentiviral Nef and Vpu Proteins.” Cell Reports 10 (4): 586–99.

67. Schmidt, Joshua M., Marc de Manuel, Tomas Marques-Bonet, Sergi Castellano, and Aida M. Andrés. 2019. “The Impact of Genetic Adaptation on Chimpanzee Subspecies Differentiation.” PLoS Genetics 15 (11): e1008485.

68. Sharp, Paul M., George M. Shaw, and Beatrice H. Hahn. 2005. “Simian Immunodeficiency Virus Infection of Chimpanzees.” Journal of Virology 79 (7): 3891–3902.

69. Silvestri, Guido, Mirko Paiardini, Ivona Pandrea, Michael M. Lederman, and Donald L. Sodora. 2007. “Understanding the Benign Nature of SIV Infection in Natural Hosts.” The Journal of Clinical Investigation 117 (11): 3148–54.

70. Snyder, Mark H., William T. London, Eveline L. Tierney, Hunein F. Maassab, and Brian R. Murphy. 1986. “Restricted Replication of a Cold-Adapted Reassortant Influenza A Virus in the Lower Respiratory Tract of Chimpanzees.” The Journal of Infectious Diseases 154 (2): 370–71.

71. Souilmi, Yassine, M. Elise Lauterbur, Ray Tobler, Christian D. Huber, Angad S. Johar, Shayli Varasteh Moradi, Wayne A. Johnston, Nevan J. Krogan, Kirill Alexandrov, and David Enard. 2021. “An Ancient Viral Epidemic Involving Host Coronavirus Interacting Genes More than 20,000 Years Ago in East Asia.” Current Biology: CB 31 (16): 3704.

72. Svardal, Hannes, Anna J. Jasinska, Cristian Apetrei, Giovanni Coppola, Yu Huang, Christopher A. Schmitt, Beatrice Jacquelin, et al. 2017. “Ancient Hybridization and Strong Adaptation to Viruses across African Vervet Monkey Populations.” Nature Genetics 49 (12): 1705–13.

73. Vetter, Michael L., and Richard T. D’Aquila. 2009. “Cytoplasmic APOBEC3G Restricts Incoming Vif-Positive Human Immunodeficiency Virus Type 1 and Increases Two-Long Terminal Repeat Circle Formation in Activated T-Helper-Subtype Cells.” Journal of Virology 83 (17): 8646–54.

74. Vingert, Benoît, Daniela Benati, Olivier Lambotte, Pierre de Truchis, Laurence Slama, Patricia Jeannin, Moran Galperin, et al. 2012. “HIV Controllers Maintain a Population of Highly Efficient Th1 Effector Cells in Contrast to Patients Treated in the Long Term.” Journal of Virology 86 (19): 10661–74.

75. Wang, Jiyao, Philippe Youkharibache, Dachuan Zhang, Christopher J. Lanczycki, Renata C. Geer, Thomas Madej, Lon Phan, et al. 2020. “iCn3D, a Web-Based 3D Viewer for Sharing 1D/2D/3D Representations of Biomolecular Structures.” Bioinformatics 36 (1): 131–35.

76. Wertheim, Joel O., and Michael Worobey. 2009. “Dating the Age of the SIV Lineages That Gave Rise to HIV-1 and HIV-2.” PLoS Computational Biology 5 (5): e1000377.

77. Wetzel, Katherine S., Yanjie Yi, Sarah T. C. Elliott, Dino Romero, Beatrice Jacquelin, Beatrice H. Hahn, Michaela Muller-Trutwin, Cristian Apetrei, Ivona Pandrea, and Ronald G. Collman. 2017. “CXCR6-Mediated Simian Immunodeficiency Virus SIVagmSab Entry into Sabaeus African Green Monkey Lymphocytes Implicates Widespread Use of Non-CCR5 Pathways in Natural Host Infections.” Journal of Virology 91 (4): e01626–16.

78. Won, Yong-Jin and Jody Hey 2004. “Divergence Population Genetics of Chimpanzees.” Molecular Biology and Evolution 22 (2): 297–307.

79. Wooding, Stephen, Anne C. Stone, Diane M. Dunn, Srinivas Mummidi, Lynn B. Jorde, Robert K. Weiss, Sunil Ahuja, and Michael J. Bamshad. 2005. “Contrasting Effects of Natural Selection on Human and Chimpanzee CC Chemokine Receptor 5.” American Journal of Human Genetics 76 (2): 291–301.

80. Worobey, Michael, Marlea Gemmel, Dirk E. Teuwen, Tamara Haselkorn, Kevin Kunstman, Michael Bunce, Jean-Jacques Muyembe, et al. 2008. “Direct Evidence of Extensive Diversity of HIV-1 in Kinshasa by 1960.” Nature 455 (7213): 661–64.

81. Worobey, Michael, Paul Telfer, Sandrine Souquière, Meredith Hunter, Clint A. Coleman, Michael J. Metzger, Patricia Reed, et al. 2010. “Island Biogeography Reveals the Deep History of SIV.” Science 329 (5998): 1487.

82. Wroblewski, Emily E., Paul J. Norman, Lisbeth A. Guethlein, Rebecca S. Rudicell, Miguel A. Ramirez, Yingying Li, Beatrice H. Hahn, Anne E. Pusey, and Peter Parham. 2015. “Signature Patterns of MHC Diversity in Three Gombe Communities of Wild Chimpanzees Reflect Fitness in Reproduction and Immune Defense against SIVcpz.” PLoS Biology 13 (5): e1002144.

83. Wroblewski, Emily E., Peter Parham, and Lisbeth A. Guethlein. 2019. “Two to Tango: Co-Evolution of Hominid Natural Killer Cell Receptors and MHC.” Frontiers in Immunology 10: 177.

84. Zhang, Yuan, Yaguang Zhang, Wangpeng Gu, and Bing Sun. 2014. “TH1/TH2 Cell Differentiation and Molecular Signals.” Advances in Experimental Medicine and Biology 841: 15–44.

