## Supplementary figures S1-S15 and tables S1, S2 for "Genetic adaptations to SIV across chimpanzee populations"

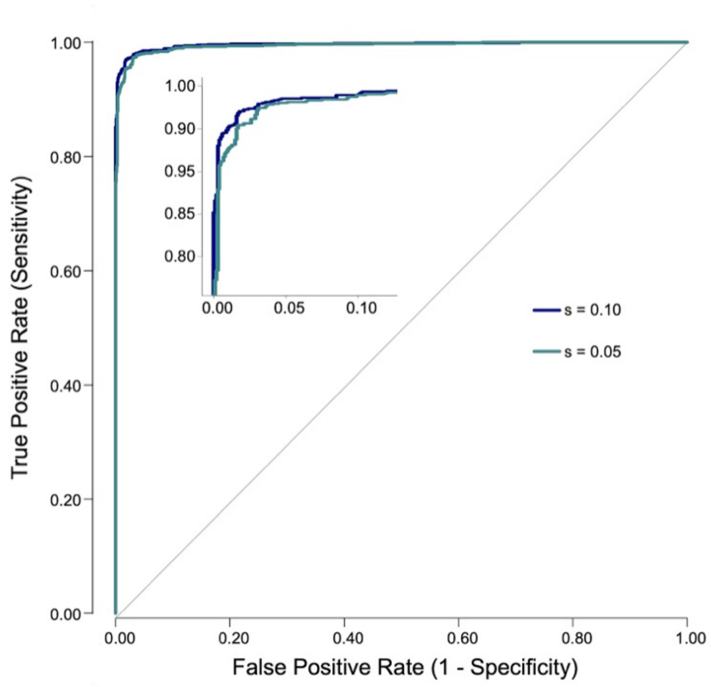


**Figure S1: Power of 3P-CLR in the ancestral central-eastern population**. Each ROC curve was generated from 1000 neutral simulations and 1000 selection simulations for each s (s=0.05, 0.1).

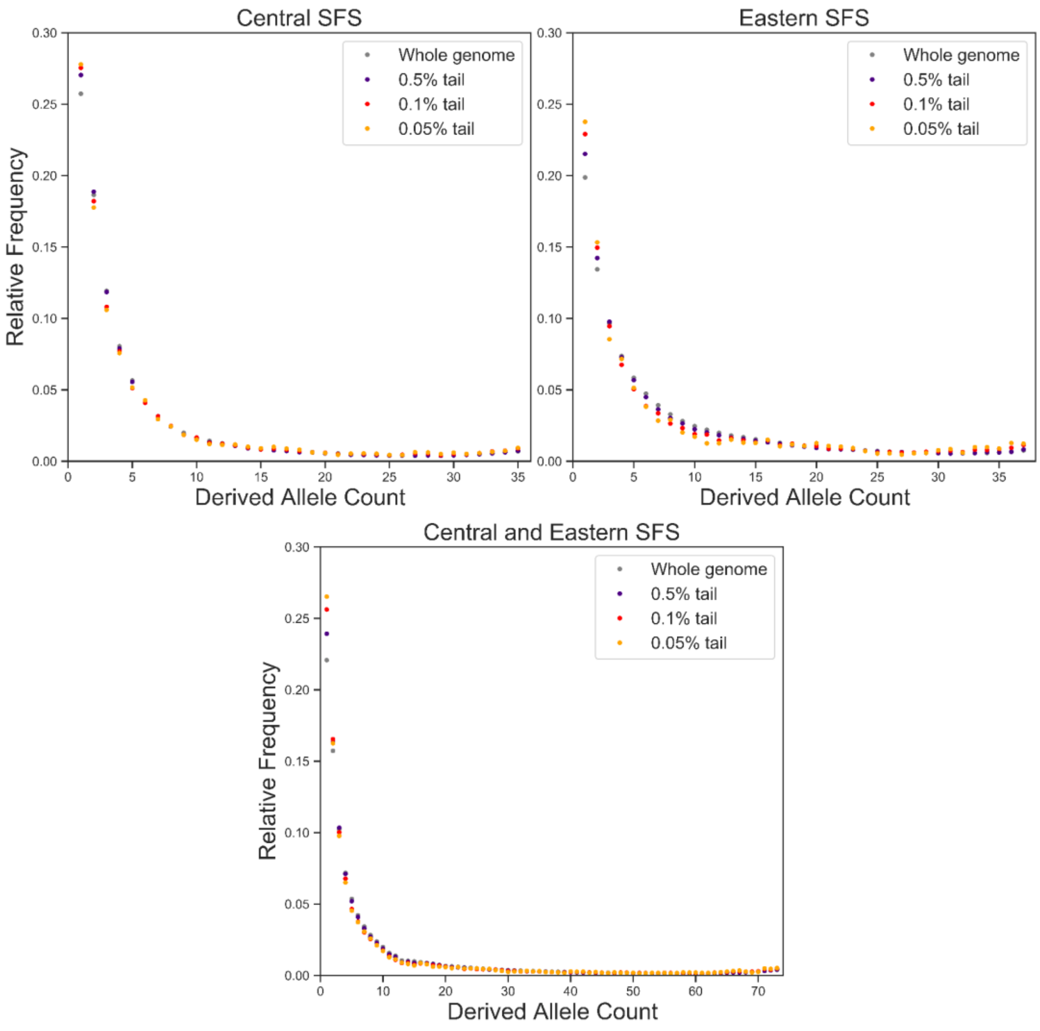


**Figure S2: Full unfolded SFS for the whole genome and each 3P-CLR tail threshold in centrals (top left), easterns (top right) and central-eastern combined (bottom).** The central-eastern combined SFS was made by simply pooling all the samples as both subspecies have nearly identical sample sizes (central: 18, eastern: 19). The SFS are all indicative of selective sweeps.

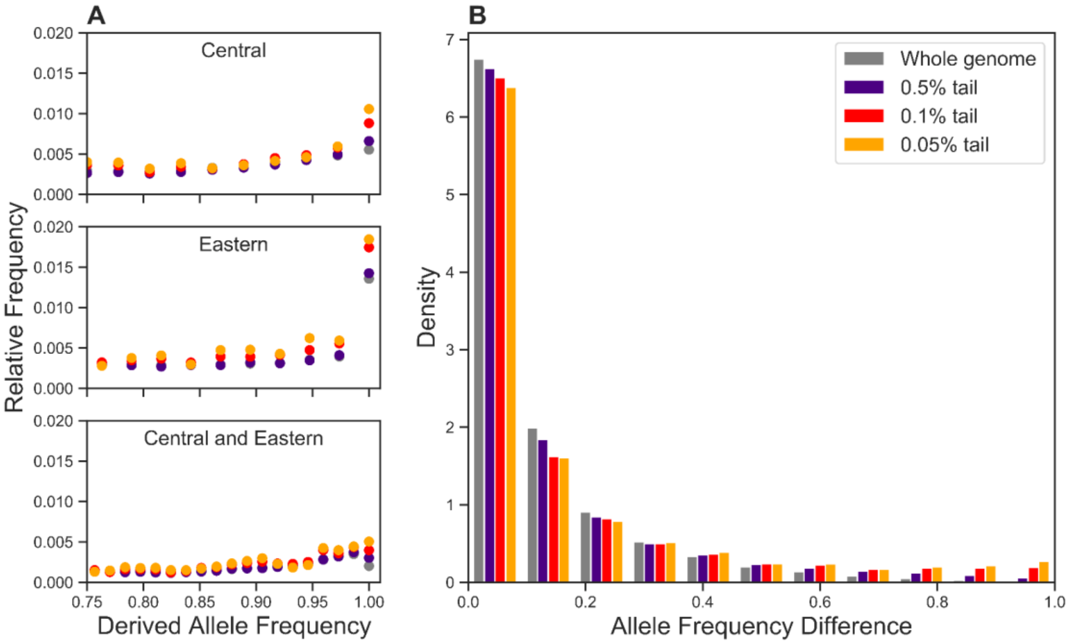


**Figure S3. Site frequency spectrum of SNPs in candidate windows.** Allele frequencies of SNPs genome-wide and at different 3P-CLR tail thresholds. A: Unfolded SFS for central, eastern and central and eastern combined. The X axis is limited to focus on high-frequency derived alleles B: Absolute DAF difference between central-eastern and Nigeria-Cameroon.

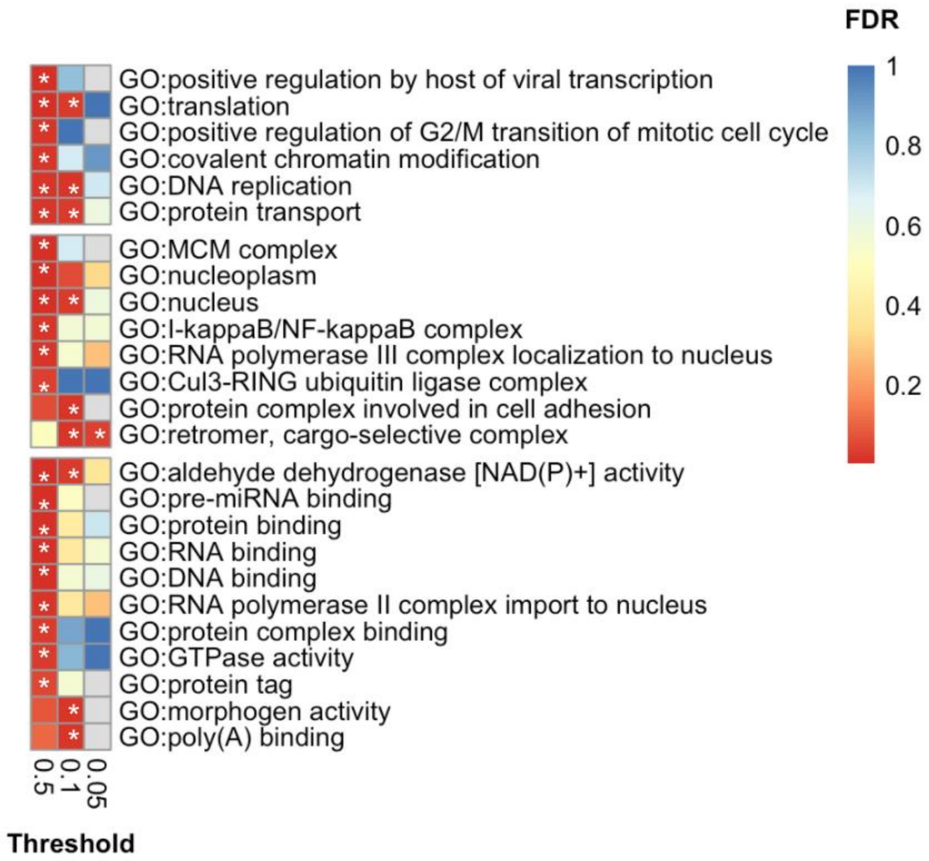


**Figure S4: Enrichment of gene ontology (GO) categories across candidate genes at different 3P-CLR quantiles in the central-eastern ancestor**. Only categories with a significant enrichment in at least one quantile (FDR<0.05) are shown. Categories are separated by GO class: Biological Process, Cellular Component and Molecular Function. Colours represent FDR values (red as highest significance). Grey represents instances where a GO category was undetected in that particular quantile. Stars indicate a significant enrichment in that 3P-CLR quantile.

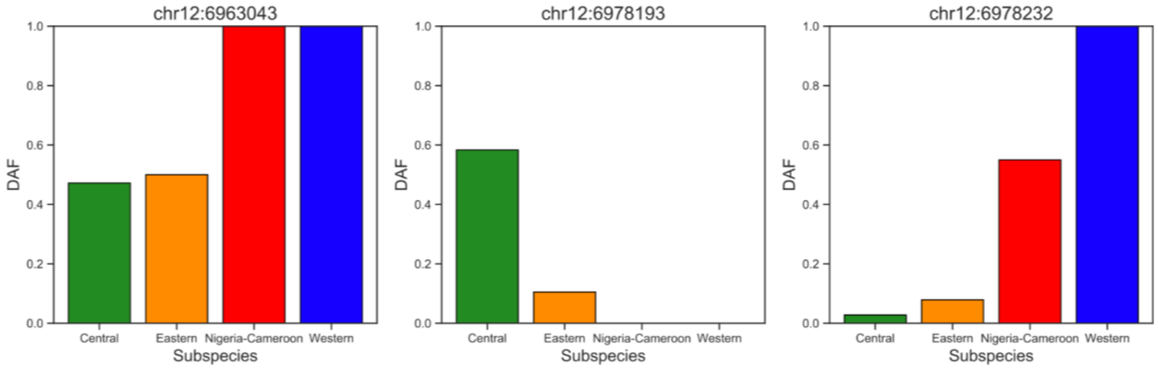


**Figure S5: DAF of the three candidate SNPs of interest in *CD4* across chimpanzee subspecies.** These candidate SNPs correspond to those highlighted in Figure 4. SNP at chr12:6963043 represents a splice variant with signatures of positive selection in the central-eastern ancestor. SNPs at chr12:6978193 (V55I SNP) and chr12:6978232 (P68T SNP) are missense variants with signatures of positive selection in centrals and the central-eastern ancestor respectively.

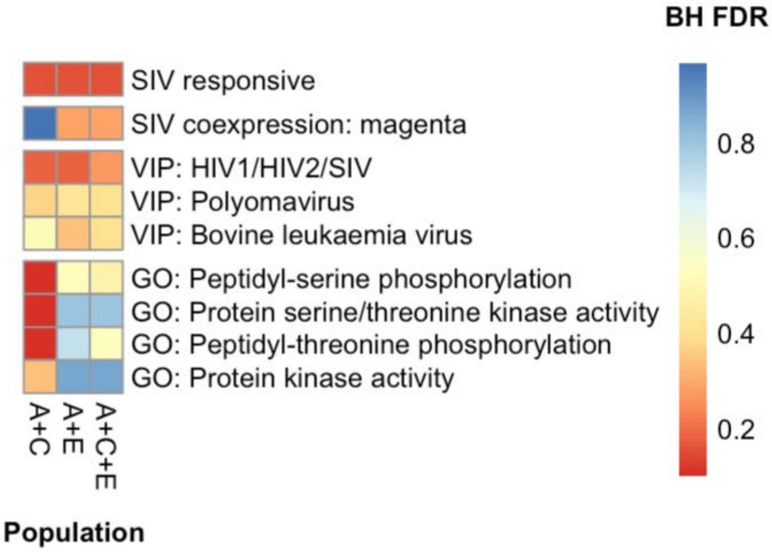


**Figure S6: Enrichment in SIV-related, VIP and GO categories of candidate targets of positive selection across populations, tested using Gowinda**. Abbreviations indicate the populations tested: central-eastern ancestor + central (A+C), central-eastern ancestor + eastern (A+E), central-eastern ancestor + central + eastern (A+C+E). Categories are separated by gene set tested: SIV responsive genes, SIV co-expression modules, VIPs and GO categories (Svardal et al. 2017; Enard et al. 2016; Enard and Petrov 2018; Ashburner et al. 2000; Jacquelin et al. 2014, 2009). Colours represent Benjamini and Hochberg corrected-FDR values (red as highest significance). We note that no category reaches significant enrichment (after correcting FDR values for the number of populations tested using a BH correction).

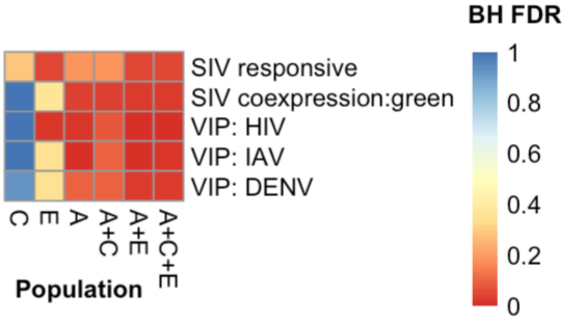


**Figure S7: Enrichment in SIV-related, and VIP categories of candidate targets of positive selection across populations, tested using *set_perm*.** Abbreviations indicate the populations tested: central (C), eastern (E), central-eastern ancestor (A), central-eastern ancestor + central (A+C), central-eastern ancestor + eastern (A+E), central-eastern ancestor + central + eastern (A+C+E). The following categories are significantly enriched in the three populations together at BH-corrected FDR < 0.05: SIV responsive genes, SIV co-expression green module, HIV/SIV

VIPs, influenza (IAV) VIPs and dengue (DENV) VIPs .All of these categories are also significantly enriched when combining the central-eastern ancestor and the eastern subspecies. The SIV co-expression green module is significantly enriched when combining the central-eastern ancestor and the central subspecies. In the single lineages, we see significant enrichment in the eastern subspecies in the SIV-responsive and HIV/SIV VIPs . While the central-eastern ancestor is significantly enriched in the SIV co-expression green module, HIV/SIV VIPs and influenza (IAV) VIPs .

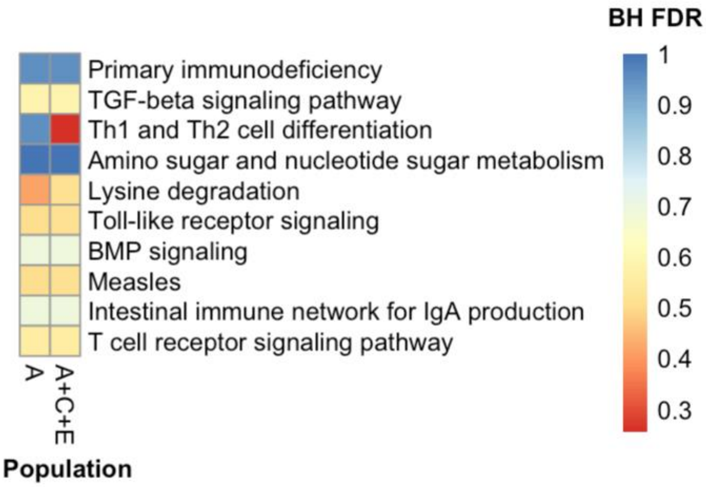


**Figure S8: Enrichment in KEGG pathways of candidate targets of positive selection across populations, tested using Gowinda**. Abbreviations indicate the population(s) tested: central-eastern ancestor (A), central-eastern ancestor + central + eastern (A+C+E). For the central-eastern ancestor we used candidate genes in the least stringent quantile (0.5), to match the number of candidates for the subspecies. Colours represent BH-corrected FDR values (red as highest significance). We note that none of the KEGG pathways reaches BH-corrected FDR < 0.05. The strongest enrichment is for the Th1 and Th2 cell differentiation pathway for A+C+E with p-value=0.00056 and BH-FDR=0.25552, which represents a nominal enrichment.

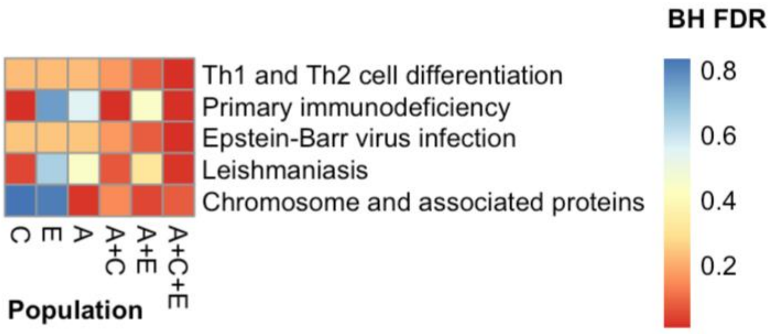


**Figure S9: Enrichment in KEGG pathways of candidate targets of positive selection across populations, tested using *set_perm*.** Abbreviations indicate the populations tested: central-eastern ancestor + central (A+C), central-eastern ancestor + eastern (A+E), central-eastern ancestor + central + eastern (A+C+E). The following KEGG categories are significantly enriched in the three populations together at BH-corrected FDR < 0.05: 'Th1 and Th2 cell differentiation', 'primary immunodeficiency', 'Epstein-Barr virus infection' and 'leishmaniasis'. The 'primary immunodeficiency' and 'leishmaniasis' categories are also significantly enriched in the central subspecies. Hence for the 'primary immunodeficiency' and 'leishmaniasis' gene sets, the central lineage is likely driving the signal when the three populations are combined. When the central-eastern ancestor and eastern subspecies are combined, we see significant enrichment in the ‘chromosome and associated proteins’ pathway, which is also significantly enriched in the central-eastern ancestor.

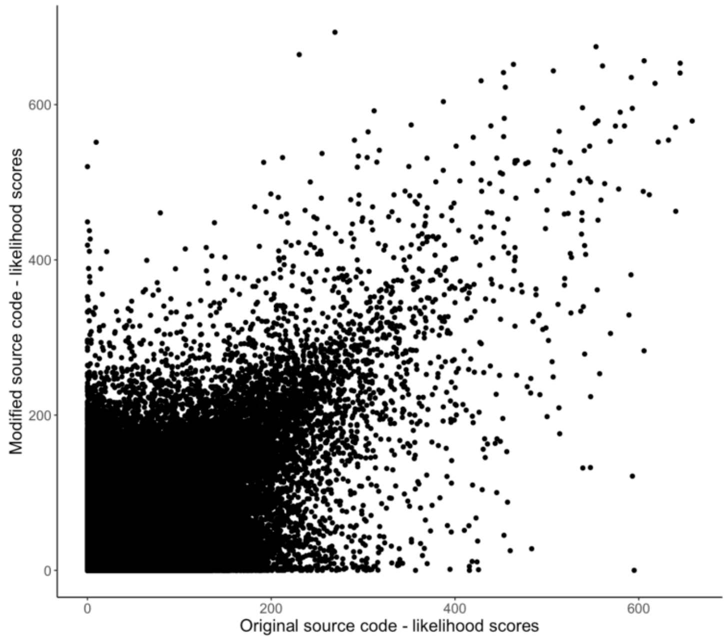


**Figure S10: Correlation between the 3P-CLR values with the original and extended code.** Likelihood scores are significantly correlated between the original and modified 3P-CLR source code (rho=0.4565, p<2.2e-16). The absence of a perfect correlation is not due to differences in the algorithm, but due to the sampling variance of SNPs in each window. Specifically, if more than 100 SNPs are present within a given window, 3P-CLR chooses 100 SNPs at random.

Thus, the same SNPs will not be sampled each time the method is run. Variation in 3P-CLR likelihood ratio scores therefore results if different SNPs were used to calculate the statistic. As expected, the correlation is weaker for low 3P-CLR but high at higher 3P-CLR values, where candidates of positive selection fall.

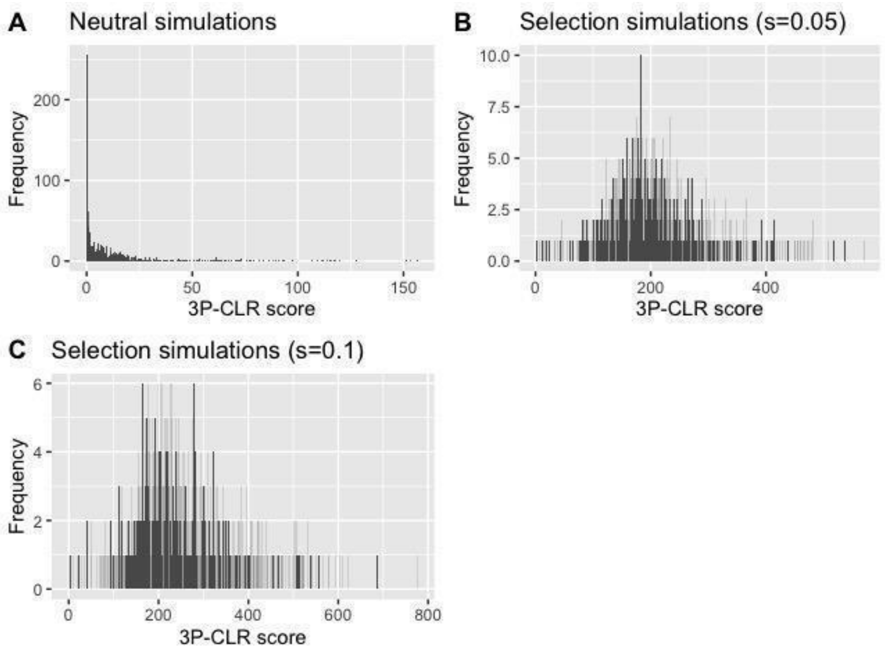


**Figure S11: Distribution of 3P-CLR scores** for data simulated under neutrality (A) and positive selection with selection coefficients of 0.05 (B) and 0.1 (C), 1000 replicates were generated in each case.

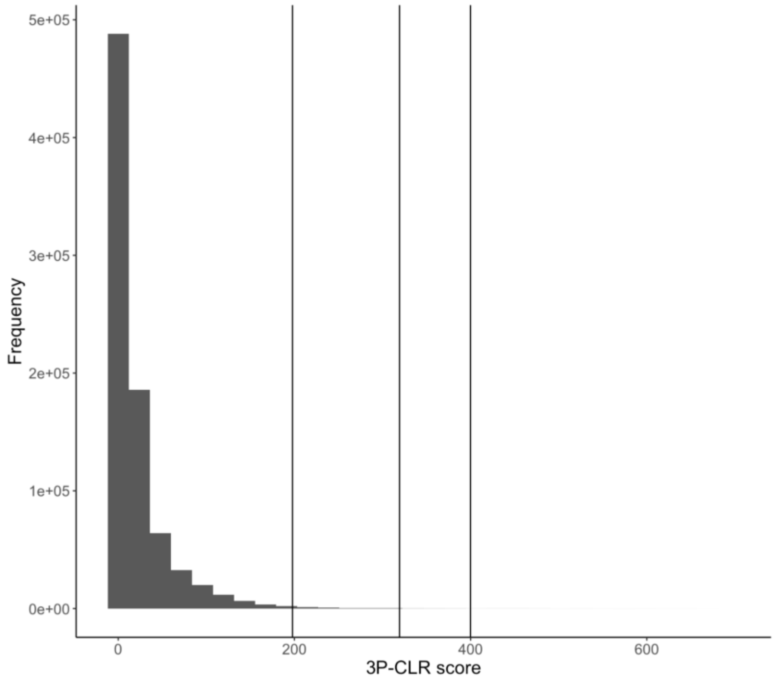


**Figure S12: Empirical distribution of 3P-CLR scores**. Vertical lines indicate the thresholds used to define candidate windows (0.5%, 0.1% and 0.05% tails of the empirical distribution).

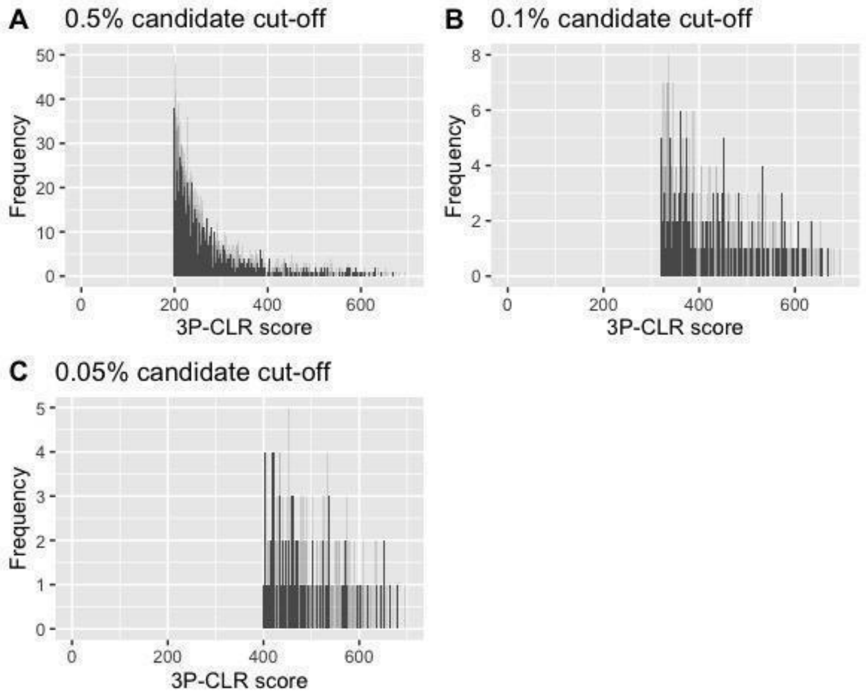


**Figure S13: Distribution of 3P-CLR scores in the tails of the empirical distribution.** (A) the 0.5%, (B) 0.1% and (C) 0.05% candidate quantiles, corresponding to 4090, 818 and 409 genomic windows respectively.

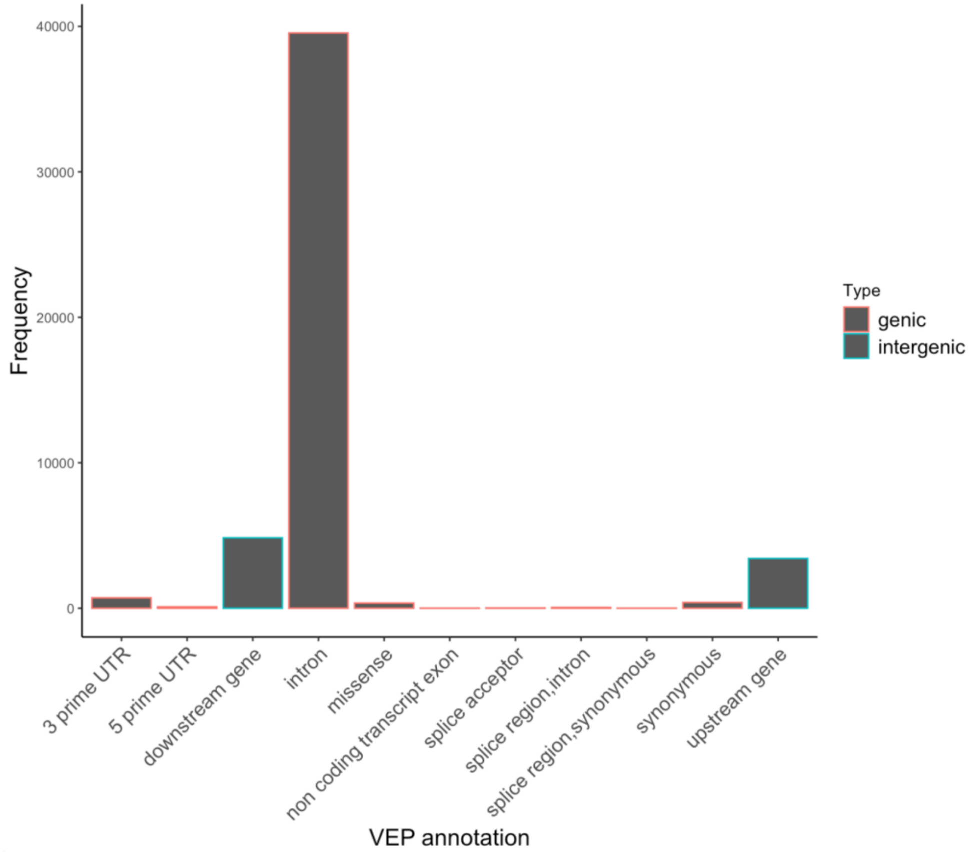


**Figure S14: Functional annotation of 20 highest PBSnj SNPs per window at the 0.5% threshold using VEP**.

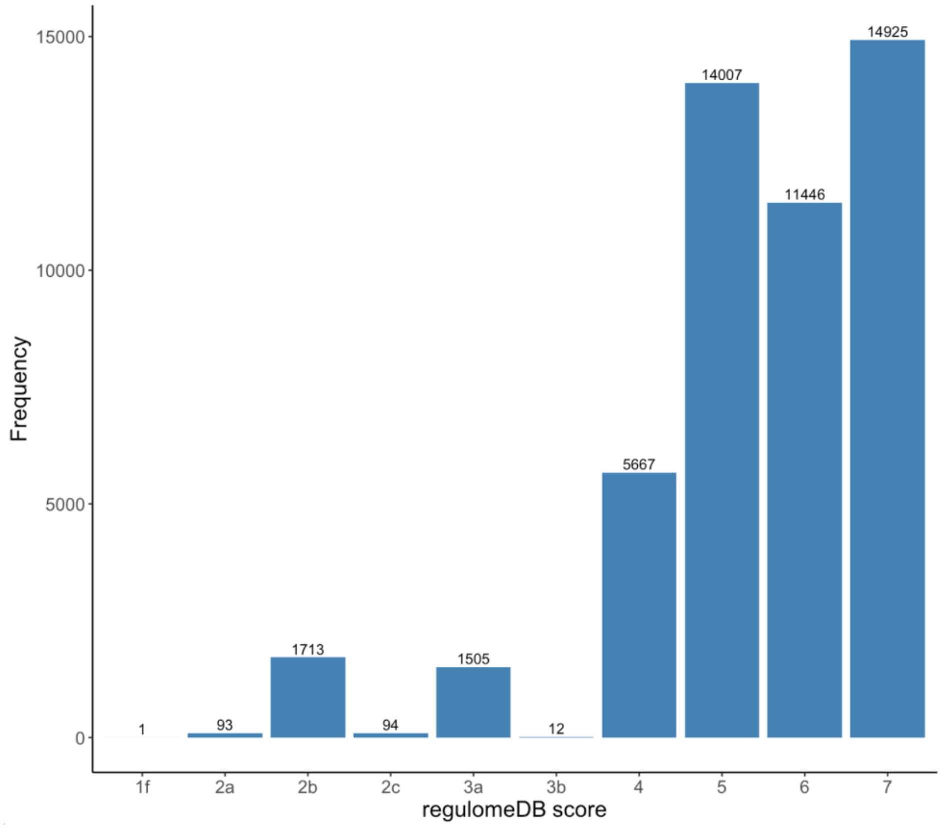


**Figure S15: Functional annotation of 20 highest PBSnj SNPs per window at the 0.5% threshold using regulomeDB**.

**Supplementary Tables**

| Number of Candidate Genes | | | | | | | |
|---|---|---|---|---|---|---|---|
| | Populations | | | | | | |
| Gene Category | C | E | A | C+E | C+A | E+A | C+E+A |
| SIV coexpression: magenta | 5 | 11 | 8 | 0 | 0 | 1 | 0 |
| SIV response | 82 | 83 | 98 | 22 | 6 | 6 | 1 |
| VIPs: HIV1/HIV2/SIV | 34 | 34 | 46 | 12 | 0 | 6 | 1 |
| Th1/Th2 differentiation pathway | 6 | 8 | 5 | 3 | 0 | 0 | 0 |

**Table S1: Number of candidate genes from each population which belong to gene categories of interest.** The number of genes which are candidates in central only, eastern only, and ancestral only are shown in the C, E and A columns respectively. The number of genes which are candidates in both central and eastern, central and ancestral, eastern and ancestral, or all three populations are shown in the C+E, C+A, E+A, and C+E+A columns respectively.

| Category | Gene | Location | | | PBSnj score | Annotation | RegulomeDB score |
|---|---|---|---|---|---|---|---|
| SIV-response | *SHCBP1* | 16 | 45562119 | 45562120 | 0.977169244867928 | upstream_gene_variant | 2b |
| SIV-response | *RAI1* | 17 | 38254584 | 38254585 | 0.979097158400808 | intron_variant | 2b |
| SIV co-expression: module green | *OGFR* | 20 | 60321593 | 60321594 | 0.540581803741123 | intron_variant | 2b |
| SIV co-expression: module green | *OGFR* | 20 | 60322673 | 60322674 | 0.515259538932669 | intron_variant | 2b |
| SIV co-expression: module green | *NUDT16L* | 16 | 4750729 | 4750730 | 0.833342835848134 | 3_prime_UTR_variant | 2b |
| SIV co-expression: module green | *NUDT16L* | 16 | 4750809 | 4750810 | 0.815198703617106 | 3_prime_UTR_variant | 2b |
| VIPs: Influenza A | *FLNC* | 7 | 130323113 | 130323114 | 0.626039709365860 | synonymous_variant(atT/atC) | 2b |
| VIPs: Influenza A | *FLNC* | 7 | 130323188 | 130323189 | 0.630704319466529 | synonymous_variant(aaT/aaC) | 2b |
| VIPs: Influenza A | *MOB2* | 11 | 1491986 | 1491987 | 0.979097158400808 | downstream_gene_variant | 2b |
| VIPs: Influenza A; HIV1/HIV2/SIV | *AKAP8L* | 19 | 15714425 | 15714426 | 0.919066385805049 | intron_variant | 2b |
| VIPs: Influenza A; HIV1/HIV2/SIV | *MYH10* | 17 | 47588373 | 47588374 | 0.869165983371999 | intron_variant | 2b |
| VIPs: Influenza A | *TNRC18* | 7 | 4030370 | 4030371 | 0.886915240089703 | intron_variant | 2b |
| VIPs: Influenza A | *TNRC18* | 7 | 4061251 | 4061252 | 0.782208048674908 | intron_variant | 2b |
| VIPs: Influenza A | *CTPS1* | 1 | 41298930 | 41298931 | 0.831830891496028 | intron_variant | 2b |
| VIPs: Influenza A | *NT5C2* | 10 | 102355543 | 102355544 | 0.794666914005296 | intron_variant | 2b |

| | | | | | | | |
|---|---|---|---|---|---|---|---|
| VIPs: Influenza A | *NT5C2* | 10 | 102355847 | 102355848 | 0.794666914005296 | intron_variant | 2a |
| VIPs: Influenza A | *HNRNPUL1* | 19 | 46464301 | 46464302 | 0.717458128517244 | intron_variant | 2b |
| VIPs: Influenza A | *HNRNPUL1* | 19 | 46469387 | 46469388 | 0.724719690820092 | intron_variant | 1f |
| VIPs: Influenza A; HIV1/HIV2/SIV | *PSMB7* | 9 | 123481626 | 123481627 | 0.762192773400031 | upstream_gene_variant | 2b |
| VIPs: Influenza A; HCMV | *RAB10* | 2A | 26388071 | 26388072 | 0.969476652360052 | 5_prime_UTR_variant | 2b |
| VIPs: HIV1/HIV2/SIV | *ITGAL* | 16 | 30634230 | 30634231 | 0.883503460161937 | intron_variant | 2b |
| VIPs: HIV1/HIV2/SIV | *SPATA5L1* | 15 | 42554379 | 42554380 | 0.833342835848134 | intron_variant | 2b |
| VIPs: HIV1/HIV2/SIV | *WHSC1* | 4 | 2021347 | 2021348 | 0.878045955284611 | 3_prime_UTR_variant | 2b |
| VIPs: HIV1/HIV2/SIV | *WHSC1* | 4 | 2021461 | 2021462 | 0.919066385805049 | 3_prime_UTR_variant | 2b |
| VIPs: HIV1/HIV2/SIV | *WHSC1* | 4 | 2021623 | 2021624 | 0.897971129460690 | 3_prime_UTR_variant | 2b |
| VIPs: HIV1/HIV2/SIV | *GNAO1* | 16 | 55330504 | 55330505 | 0.639189125279027 | intron_variant | 2b |
| VIPs: HIV1/HIV2/SIV | *SPATS2* | 12 | 39874385 | 39874386 | 0.823158026244118 | intron_variant | 2b |
| VIPs: HIV1/HIV2/SIV | *SMG6* | 17 | 2083071 | 2083072 | 0.904042285992019 | intron_variant | 2a |
| VIPs: HIV1/HIV2/SIV | *SMG6* | 17 | 2083141 | 2083142 | 0.904042285992019 | intron_variant | 2b |
| VIPs: HIV1/HIV2/SIV | *SMG6* | 17 | 2083347 | 2083348 | 0.904042285992019 | intron_variant | 2b |
| VIPs: HIV1/HIV2/SIV | *RAD23A* | 19 | 13216455 | 13216456 | 0.958872156258472 | upstream_gene_variant | 2b |
| VIPs: HIV1/HIV2/SIV | *CERS2* | 1 | 129153708 | 129153709 | 0.777225715621888 | intron_variant | 2b |
| VIPs: HCMV | *USP54* | 10 | 72098548 | 72098549 | 0.477321510964906 | missense_variant (T/A Aca/Gca) | - |

| | | | | | | | |
|---|---|---|---|---|---|---|---|
| VIPs: ADV | *RAB34* | 17 | 28361600 | 28361601 | 0.712341200918947 | upstream_gene_variant | 2b |
| VIPs: ADV | *RAB34* | 17 | 28361637 | 28361638 | 0.698462506757592 | upstream_gene_variant | 2b |
| VIPs: ADV | *RAB34* | 17 | 28362090 | 28362091 | 0.724251911659941 | synonymous_variant(ccG/ccC) | 2b |

**Table S2: SNPs of interest in genes identified within candidate windows of positive selection at the 0.5% threshold in the central-eastern ancestor, additional to *CD4.*** SNPs with regulomeDB scores of 1f, 2a, 2b, 2c are considered to have putative significant regulatory function, due to the lack of eQTL data for chimpanzees. Category indicates the gene set each gene belongs to. Abbreviations: HCMV and ADV indicate human cytomegalovirus and adenovirus respectively.
