## Supplementary note of set_perm method for "Genetic adaptations to SIV across chimpanzee populations"

### *set_perm* SOM note

Joshua Schmidt

To perform gene set enrichment tests on more than one lineage I developed ***set_perm***, which implements a permutation-based enrichment test that for a single lineage is equivalent to Gowinda (R. Kofler and Schlotterer 2012) but that accounts for genes with signatures of natural selection in more than one lineage. *set_perm* is written in python and is dependent on numpy, pandas and pytables. The code is available at https://github.com/joshuamschmidt/set_perm.

As with Gowinda, a permutation set is derived by random sampling from a background set of SNPs (or windows). The genes intersecting the sampled SNPs then form a permutation set. *set_perm* follows the ‘gene mode’ setting of Gowinda, wherein a permutation set contains the same number of genes as the observed set, and a gene is only counted once even if multiple SNPs intersecting that gene are sampled.

A joint test of lineages is performed by generating independent permutation sets for each lineage and combining them across lineages into a joint set. The total observed number of genes is thus the sum of the observed number of genes across lineages. The p-values and FDR-corrected significance are calculated using permutation as described for Gowinda (R. Kofler and Schlotterer 2012), and thus corrects for gene length bias and clustering of paralogs. In this case, or each gene set (S) and permutation (i), the joint permutation set n is obtained by summing the number of genes (n) per lineage (k) i.e.

$$joint_{Si} = \sum_{1}^{k} n_k$$

The total observed number of genes is the sum of the observed number of genes in the different lineages. This means that the same gene can be counted more than once if it is a candidate in more than one lineage.

**Comparison to Gowinda.**

To validate the performance of *set_perm* against the established method Gowinda (Kofler and Schlötterer 2012) when testing for gene set enrichment in one lineage, we tested for enrichment of the 0.5% candidate windows of the central-eastern chimpanzee ancestor in KEGG categories (Qiu 2013), determined with 100,000 permutations (Figure 1).

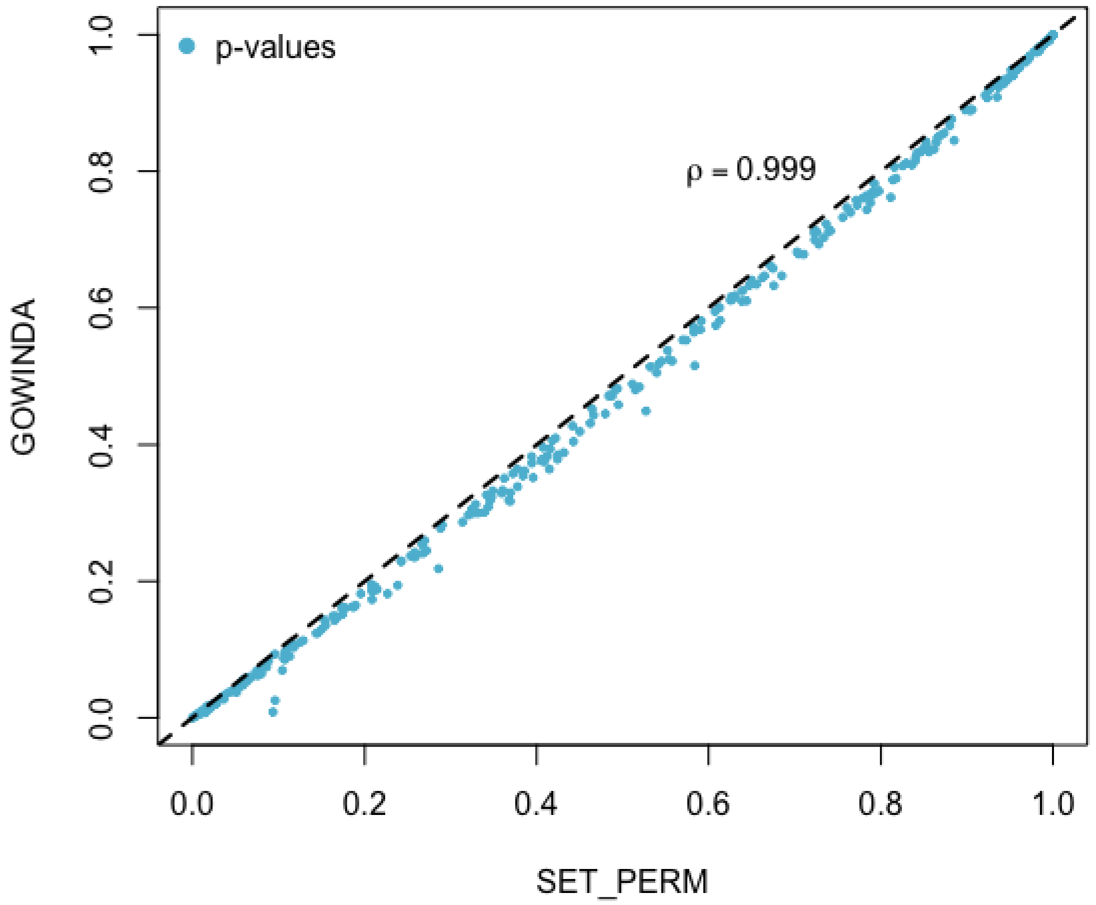


**Figure 1: Comparison of empirical p-values generated by *set_perm* and Gowinda.** X and Y axes: empirical p-values for KEGG gene set enrichment tests.

Figure 1 shows a very high concordance of empirically determined p-values calculated under *set_perm* and Gowinda with a Spearman rho = 0.999 (p-value < 2.2e-16). Notably, the trend is for p-values to be below identity, suggesting that *set_perm* is slightly more conservative in calculating empirical p-values than Gowinda. This result confirms the functional equivalence of basic permutation p-value estimation procedures in *set_perm* and Gowinda, including correcting for gene length bias and clustering of paralogs. Hence when testing for gene set enrichment in one lineage the performance of Gowinda and *set_perm* are equivalent.

### Joint lineage tests: robustness to false positives

The joint test procedure in *set_perm* sums the number of genes per functional category per lineage, for both the observed and permutation sets. As an example, for a gene category with total size of 30 genes, and with 10 candidates in each of two lineages, the joint n candidates is simply 10 + 10 = 20. Analogously, the joint gene category size is now 30 + 30 = 60. Of note, a gene which is in both candidate sets (ie. a candidate in two lineages) contributes twice. Hence the union of candidate genes <= number of joint candidates. This enables repeated adaptation to be statistically tested in the enrichment hypothesis, and hence marks an extension to the functionality of Gowinda.

We first confirm that in the absence of enrichment, a joint test of random sets of candidates does not generate a false signal of gene enrichment. We generated two candidate sets by random sampling from the central-eastern ancestor background SNPs. Note that two random sets may have genes in common. Each set consisted of 2000 SNPs. Again, KEGG pathways were used to define the functional categories for enrichment testing. This analysis confirms that results are again highly congruent between *set_perm* and Gowinda, and neither candidate set is enriched for any KEGG category (Figure 2).

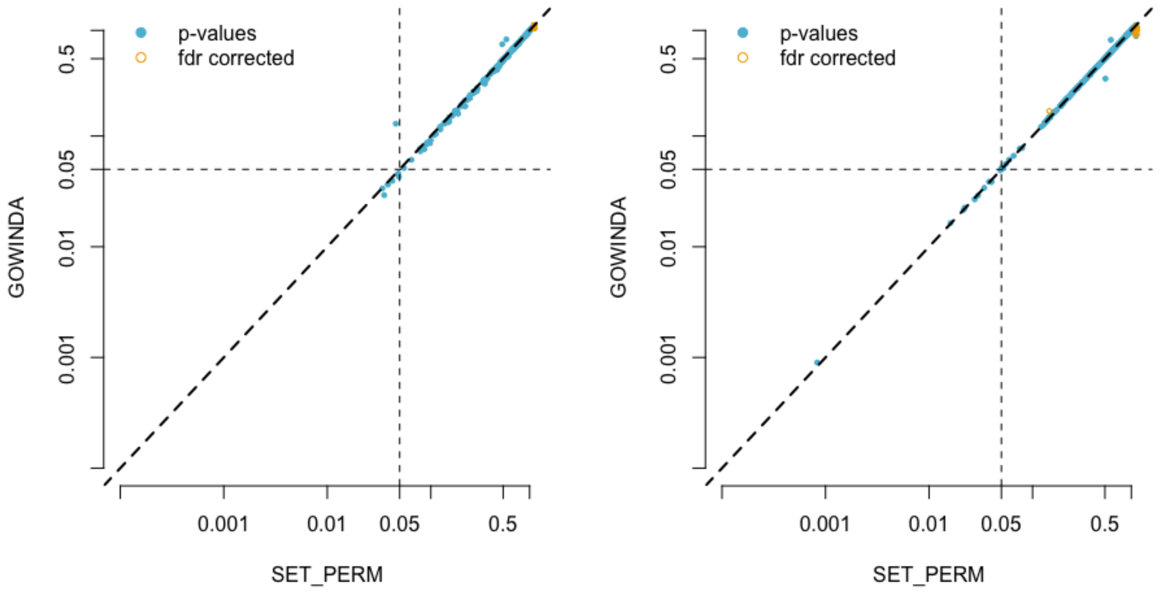


**Figure 2: Random candidate SNPs show no significant enrichments.**
Both *set_perm* and Gowinda successfully control for false positives at an FDR corrected p-value threshold of 0.05, for two independent candidate sets generated by random sampling of 2000 central-eastern ancestor background SNPs. X and Y axes: log10 transformed p-values.

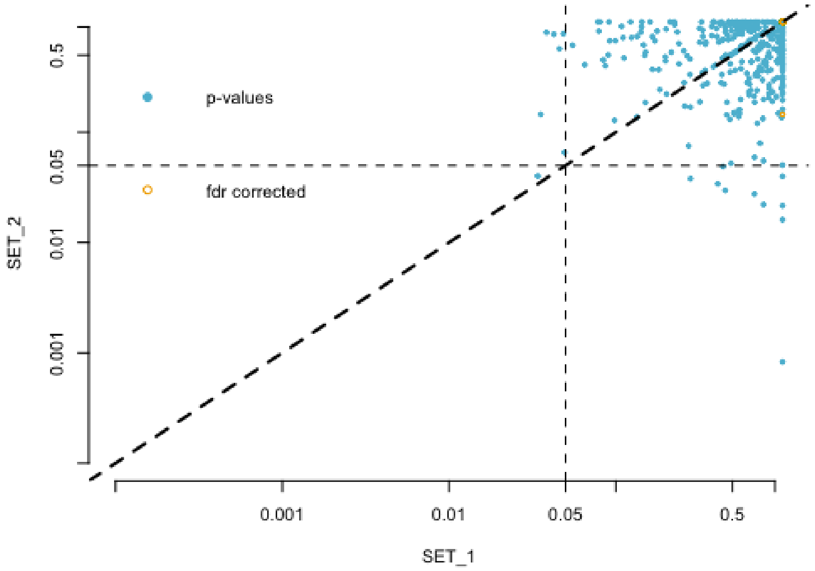
p-values
fdr corrected
SET_2
0.5
0.05
0.01
0.001
0.001
0.01
0.05
0.5
SET_1

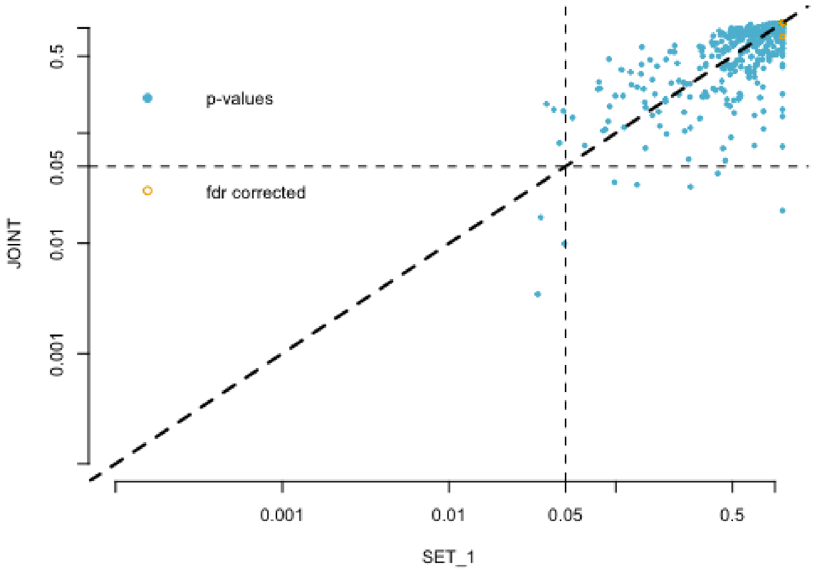
p-values
fdr corrected
JOINT
0.5
0.05
0.01
0.001
0.001
0.01
0.05
0.5
SET_1

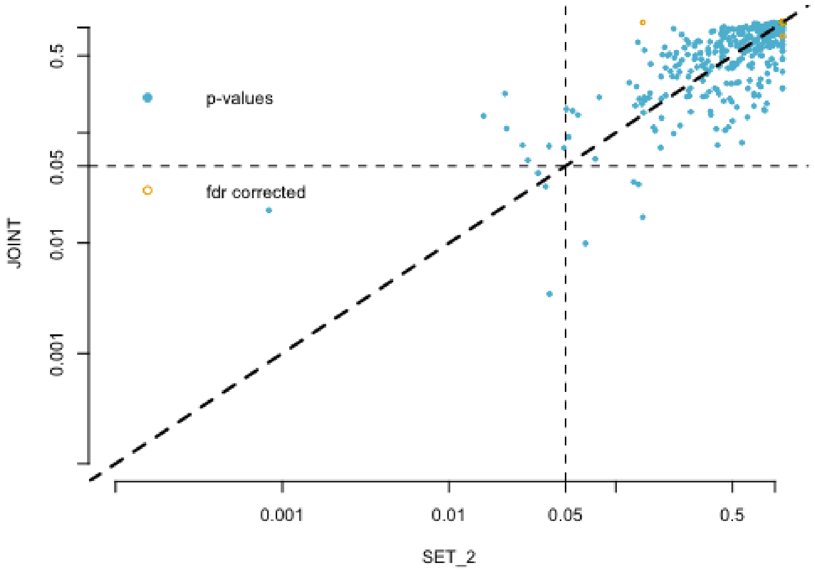
p-values
fdr corrected
JOINT
0.5
0.05
0.01
0.001
0.001
0.01
0.05
0.5
SET_2

**Figure 3: *set_perm* joint lineage test is robust to false positives.**
Top row: Plotting enrichment p-values calculated under *set_perm* for each of the two independent candidate sets (obtained by random sampling above) against each other, shows that as expected these two random candidate sets are largely uncorrelated and do not exhibit significant enrichments individually.
Middle and bottom rows: Plotting the joint test p-values against either candidate set shows an increase in the correlation, as expected because both SET1 and SET 2 are part of the joint set, but there remains no significant categories, indicating adequate FDR control.
All: x and y axes: log10 transformed p-values.

As the joint test is the sum of the number of candidate genes per functional category, the p-values of the joint test and each of the individual candidate sets are correlated –more so than the p-values of the individual candidate sets are (Figure 3, left column). However, the joint test shows no significant FDR corrected p-values (Figure 3, middle and bottom columns). This indicates that the joint test itself does not result in false positive enrichment tests.

#### Joint lineage tests: ability to detect enriched categories

Finally, to show that a joint test increases power to find gene set enrichment in the presence of targets of selection in multiple lineages, we generated two further candidate SNP lists: the random lists presented in Figure 2, with the artificial addition of 2-3 SNPs in candidate genes from the four KEGG categories with size 5 < n < 10 (hsa00072, hsa00232, hsa00524, hsa04122). This is analogous to the Gowinda validation procedure, except that here candidate SNPs are split between the two lineages.

As shown in Figure 4 (left column), there is no significant KEGG category in either of the candidate sets after FDR correction. However, the joint test shows significant enrichment for 3 of 4 of the manipulated categories. The fourth, hsa00542 has FDR corrected p-value of 0.15. This strongly indicates that *set_perm* can leverage the information obtained from selection statistics in two or more lineages to find evidence of enrichment for function.

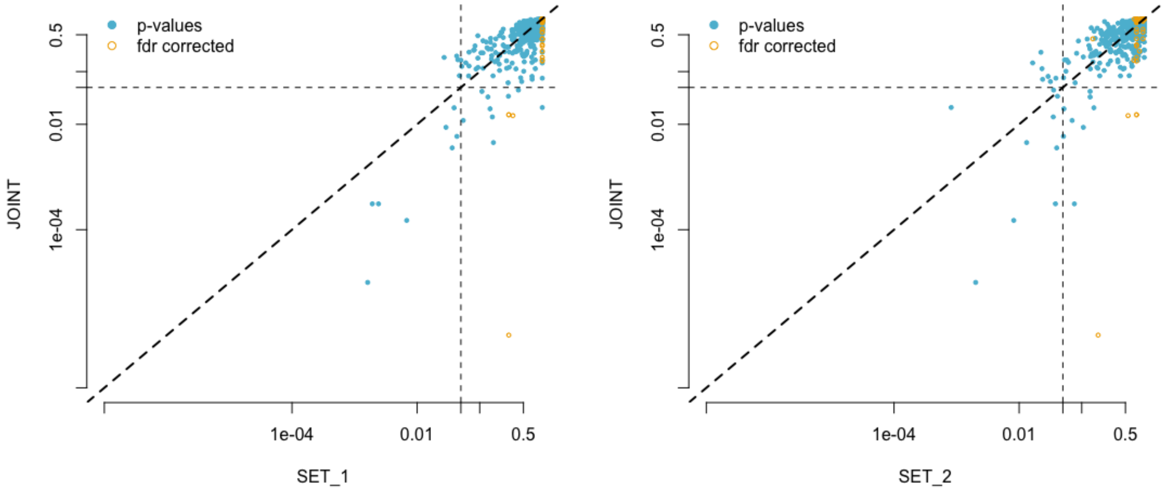


**Figure 4: set_perm joint lineage test increases power.**
Neither candidate set 1 (x-axis, left panel) nor candidate set 2 (x-axis, right panel) exhibit significant enrichment in target KEGG gene sets, when candidate SNPs in target genes are split between taxa. By combining candidates across taxa, significant enrichments are now discovered for 3 of 4 targeted KEGG sets (y-axis, both panels).
Both: x and y axes: log10 transformed p-values.

#### References

Kofler, Robert, and Christian Schlötterer. 2012. “Gowinda: Unbiased Analysis of Gene Set Enrichment for Genome-Wide Association Studies.” *Bioinformatics* 28 (15): 2084–85.

Qiu, Yu-Qing. 2013. “KEGG Pathway Database.” *Encyclopedia of Systems Biology*. https://doi.org/10.1007/978-1-4419-9863-7_472.
